# *Cnot3* is required for male germ cell development and spermatogonial stem cell maintenance

**DOI:** 10.1101/2023.10.13.562256

**Authors:** Qing Chen, Safia Malki, Xiaojiang Xu, Brian Bennett, Brad L. Lackford, Oleksandr Kirsanov, Christopher B. Geyer, Guang Hu

## Abstract

The foundation of spermatogenesis and lifelong fertility is provided by spermatogonial stem cells (SSCs). SSCs divide asymmetrically to either replenish their numbers (self-renewal) or produce undifferentiated progenitors that proliferate before committing to differentiation. However, regulatory mechanisms governing SSC maintenance are poorly understood. Here, we show that the CCR4-NOT mRNA deadenylase complex subunit CNOT3 plays a critical role in maintaining spermatogonial populations in mice. *Cnot3* is highly expressed in undifferentiated spermatogonia, and its deletion in spermatogonia resulted in germ cell loss and infertility. Single cell analyses revealed that *Cnot3* deletion led to the de-repression of transcripts encoding factors involved in spermatogonial differentiation, including those in the glutathione redox pathway that are critical for SSC maintenance. Together, our study reveals that CNOT3 – likely via the CCR4-NOT complex – actively degrades transcripts encoding differentiation factors to sustain the spermatogonial pool and ensure the progression of spermatogenesis, highlighting the importance of CCR4-NOT-mediated post-transcriptional gene regulation during male germ cell development.

**Highlights:**

- *Cnot3* is predominantly expressed in undifferentiated spermatogonia, and its expression decreases in later stages.
- Conditional deletion of *Cnot3* leads to total germ cell loss and male infertility.
- CNOT3 represses transcripts encoding factors involved in spermatogonial differentiation.
- CNOT3 protects spermatogonial stem cells maintenance via GSH/redox pathway.

**Graphical Abstract:** Cnot3 promotes SSC self-renewal by balancing ROS production via the regulation of GSH production

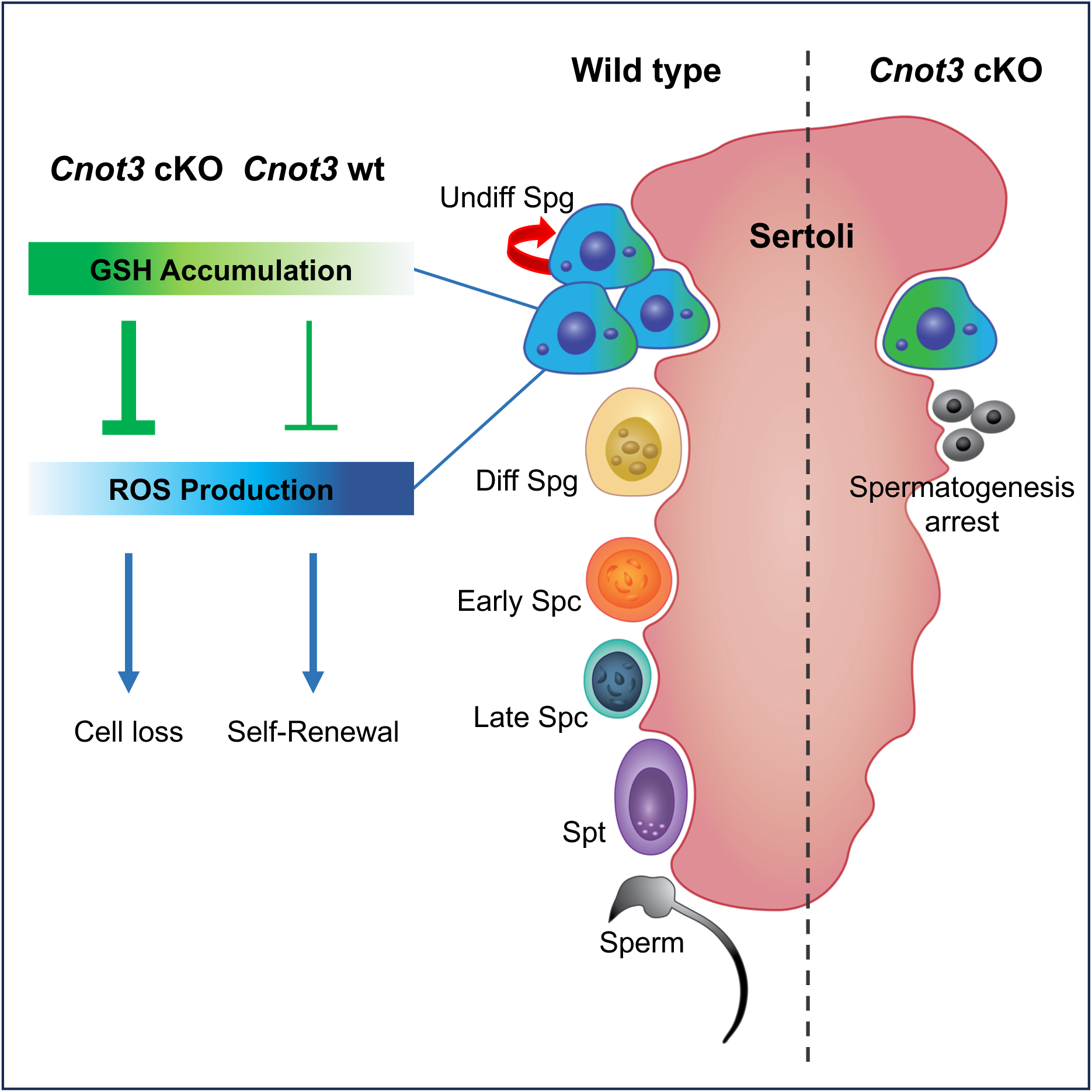

## Introduction

Spermatogenesis is a stem cell-based developmental program that produces large numbers of haploid gametes every day to sustain lifelong male fertility. Originating in the fetal testis, prospermatogonia (also called gonocytes), transition after birth into spermatogonia, a subset of which are spermatogonial stem cells (SSCs) (de Rooij, 2001). These SSCs balance their divisions during steady-state spermatogenesis to both replenish their population (via self-renewal) and produce undifferentiated progenitors that proliferate before differentiating (Oatley and Brinster, 2006). Differentiation occurs in response to retinoic acid (RA), which ensures an irreversible commitment to ultimately enter meiosis (Gewiss et al., 2021; Griswold, 2016; Koubova et al., 2006; Velte et al., 2019). After five divisions, these differentiating spermatogonia become preleptotene spermatocytes, which divide one final time before entering meiosis. Following the lengthy meiotic program, spermatocytes divide twice to form haploid round spermatids that undergo the morphogenetic program of spermiogenesis to become sperm (de Kretser, 1998; de Rooij and Russell, 2000).

The maintenance of the SSC pool is orchestrated by both transcriptional and posttranscriptional mechanisms (Legrand and Hobbs, 2018). While the roles of several RNA binding proteins (RBPs) have been described during spermatogenesis, only a handful have been implicated in SSCs such as NANOS2, DND1, and TRIM71 (Du et al., 2020; Legrand and Hobbs, 2018; Niimi et al., 2019; Zhou et al., 2015).

We previously reported that the CNOT1, CNOT2, and CNOT3 subunits of the CCR4-NOT complex maintain the pluripotent state in mouse embryonic stem cells (ESCs) by preventing their differentiation into extraembryonic lineages (Zheng et al., 2012). Deletion of *Cnot3* in ESCs reveals a role in promoting the deadenylation and subsequent degradation of mRNAs encoding differentiation factors, revealing a requirement for mRNA decay in the maintenance of the pluripotent state (Hu et al., 2009; Zheng *et al*., 2012). In mice, deletion of *Cnot3* causes embryonic lethality at the blastocyst stage due to loss of the inner cell mass (Neely et al., 2010).

In this study, we generated male germ cell-specific *Cnot3* knockout (KO) mice and identified a critical role for CNOT3 in maintaining the SSC reserve. We show that in the developing testis of the *Cnot3* KO animals, SSC and progenitor populations in the spermatogonia were depleted, leading to loss of the germline and male sterility. Furthermore, *Cnot3* deletion resulted in the de-repression of transcripts encoding factors involved in SSC differentiation, including those in the glutathione redox pathway that are critical for SSC maintenance. Together, our results indicate that CNOT3-mediated mRNA decay serves as an essential post-transcriptional regulatory mechanism in SSC maintenance.

## Results

### *Cnot3* deletion in adult mice results in germ cell loss and male infertility

As a first step to explore the role of CNOT3 during mouse spermatogenesis, we assessed *Cnot3* transcript levels in published scRNA-seq data (Green et al., 2018). We found that *Cnot3* mRNAs are abundant in spermatogonia, but later decrease in spermatocytes, round spermatids, and elongating spermatids (Suppl Fig 1A). At the protein level, immunostaining results showed that CNOT3 is detectable in ZBTB16 (also known as PLZF) positive cells located along the basement membrane of seminiferous tubules in adult mouse testes (Suppl Fig 1B-C). ZBTB16-positive cells in the testes are undifferentiated spermatogonia, comprising both the SSC and progenitor populations (Buaas et al., 2004; Costoya et al., 2004; Hobbs et al., 2010). Thus, we posited a role for CNOT3 in undifferentiated spermatogonia.

To elucidate the function of CNOT3 in spermatogonia, we generated a conditional knockout mouse model in which *Cnot3* was specifically deleted in germ cells by the tamoxifen-inducible *Ddx4*-Cre transgene (*Cnot3*^flox/flox^; *Ddx4*-creER) (Zheng et al., 2016), hereafter referred to as *Cnot3*-cKO. To induce Cre expression, we dosed mice with tamoxifen (4-OHT) at 4-weeks of age and examined the fertility of adult (8-weeks old) *Cnot3*-cKO males (Fig1 A). We first confirmed *Cnot3* deletion by immunostaining (Fig 1B). We then performed a breeding study for six weeks by pairing 8-weeks old wild-type (WT) control and *Cnot3*-cKO males with WT females (Fig 1A). While control males produced 1-2 litters, all but one *Cnot3*-cKO male were sterile and had no litters (Fig 1C). We next euthanized additional tamoxifen-treated *Cnot3*-cKO males at 6- and 8-weeks old and recorded body, testis, and epididymis weights. Although *Cnot3*-cKO body weights did not differ from controls, we observed a significant reduction in the ratios of testis and epididymis to body weights (Fig 1D). Histological analyses revealed *Cnot3*-cKO males had smaller testes (Fig 1E), with multinucleated degenerating cells in testes of 6-week old *Cnot3*-cKO mice and an absence of germ cells in seminiferous tubules of 8-week old mice (Fig 1E). The absence of germ cells was confirmed by immunostaining, in which the pan germ cell-specific marker TRA98 was nearly undetectable in *Cnot3*-cKO testes (Fig 1F). In contrast, the Sertoli cell-specific marker SOX9 showed similar expression pattern in WT and cKO testis, indicating that Sertoli cell numbers were not altered in cKO mice (Fig 1F). The infertility and total loss of germ cells in *Cnot3*-cKO testes suggested the loss of SSCs and thereby a potential role of *Cnot3* in SSC maintenance.

**Figure 1.**
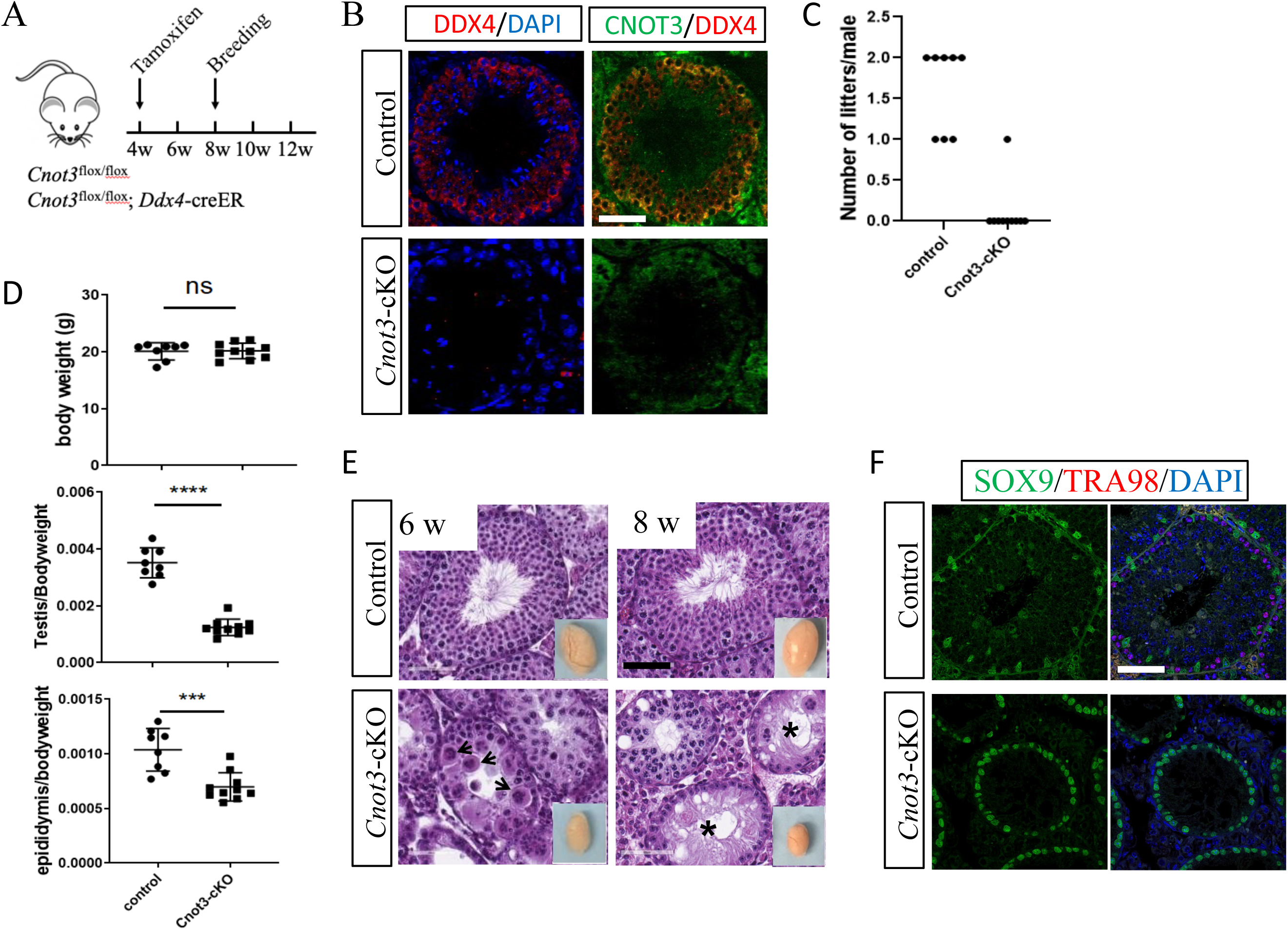
*Cnot3* deletion results in germ cell loss and male infertility. (A) Schematic of conditional deletion of *Cnot3* in male germ cell upon Tamoxifen administration. (B) Merged channels of single confocal sections of 8-weeks old Control and Cnot3-cKO testis showing CNOT3 (green) and DDX4 (red) expression with DAPI (blue). (C) Quantification of alive litters obtained from WT female mated with Control and *Cnot3*-cKO males. (D) Quantification of body weight in grams, testis to body weight ratio and, epididymis to body ratio weight of 8 weeks-old control and *Cnot3*-cKO males (two-tailed unpaired Student’s t test; (ns) for p > 0.05, (^∗∗∗^) for p < 0.001, and (^∗∗∗^) for p < 0.0001). (E) Hematoxylin and eosin-stained sections of testis of 6 and 8 weeks-old control and *Cnot3*-cKO males. Insert shows whole testis. (F) Merged channels of single confocal sections of 8-weeks old Control and *Cnot3*-cKO testis showing SOX9 (green) and TRA98 (red) expression with DAPI (blue).

### *Cnot3* deletion in neonatal mice results in spermatogonia pool depletion

During normal development, by postnatal day-3 (P3) quiescent prospermatogonia reenter the cell cycle, migrate to the basement membrane of the testis cords, and transition into spermatogonia (Drumond et al., 2011; Nagano et al., 2000). This foundational population of spermatogonia contains both precursors that will form future adult SSCs, as well as undifferentiated progenitors and differentiating spermatogonia that will initiate the first wave of spermatogenesis (Hermann et al., 2015; Kluin and de Rooij, 1981; Law et al., 2019; Nikolova et al., 1997; Yoshida et al., 2006). Based on this, we can examine the role of *Cnot3* in SSCs more specifically by deleting *Cnot3* in this newly formed spermatogonial population in neonatal testes. We dosed *Cnot3*^flox/flox^;*Ddx4-* CreER mice and *Cnot3*^flox/flox^ control littermates with tamoxifen from P1 to P3 and collected testes at P6, P10, P14, and P21 (Fig 2A). At these ages, testes contain the following most advanced germ cell types: P6 = spermatogonia, P10 = leptotene spermatocytes, P14 = pachytene spermatocytes, and P21 = round spermatids. We first immunostained testes for CNOT3, the pan germ cell marker DDX4, and the Sertoli cell marker SOX9. We observed absence of CNOT3 protein by P6 (Fig 2B-C). At P14, seminiferous tubules of *Cnot3*-cKO mice contained very few DDX4+ germ cells, but numerous SOX9+ Sertoli cells (Fig 2B, 2F and Suppl Fig 2). Consistent with that, *Cnot3*-cKO testes appeared smaller by P14, and the histology analysis showed that while the newly-formed seminiferous tubule structures appeared intact, their diameters were decreased, and the interstitial space was increased (Fig 2D). Average testis-to-body weight ratios of these *Cnot3*-cKO P14 testes were significantly reduced as compared to control testes (Fig 2E). These observations indicated that CNOT3 deletion in the spermatogonia of newborn animals caused complete germ cell loss, similar to results in *Cnot3*-cKO adult mice.

**Figure 2.**
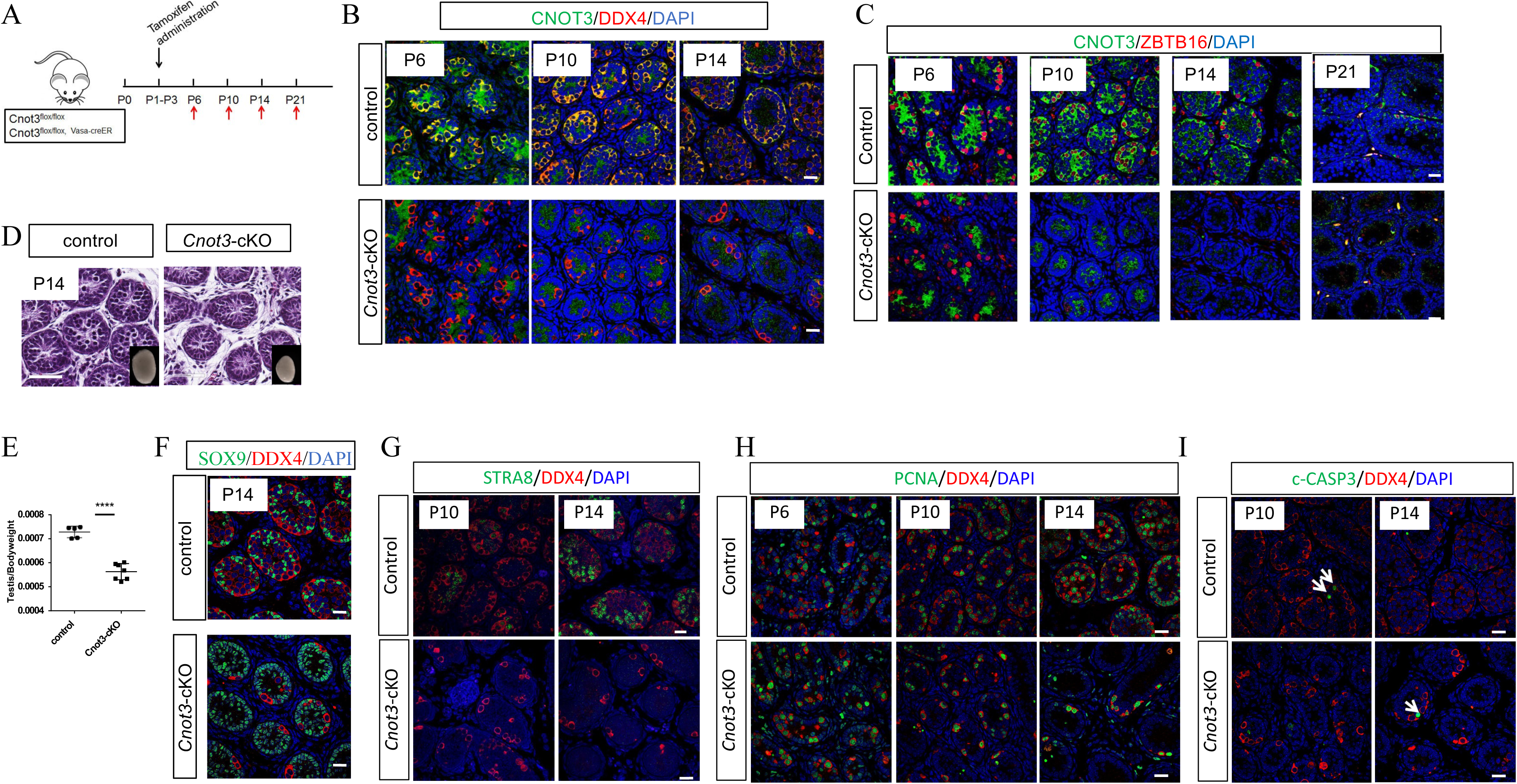
*Cnot3* deletion causes depletion of ZBTB16-spermatogonia and decrease of SSC maintenance. (A) Experimental scheme to achieve and analyze the deletion of *Cnot3* in neonatal testis. (B) Merged channels of single confocal sections of P6, P10 and P14 Control and *Cnot3*-cKO testis showing CNOT3 (green) and DDX4 (red) expression with DAPI (blue). (C) Merged channels of single confocal sections of P6, P10, P14 and P21 Control and *Cnot3*-cKO testis showing CNOT3 (green) and ZBTB16 (red) expressing cells with DAPI (blue). (D) Hematoxylin and eosin-stained sections of testis of P14 WT and *Cnot3*-cKO males. (E) Quantification of testis to body weight ratio of P14 control and *Cnot3*-cKO males (two-tailed unpaired Student’s t test; (^∗∗∗^) for p < 0.0001). (F) Merged channels of single confocal sections of P14 Control and *Cnot3*-cKO testis showing SOX9 (green) and DDX4 (red) expressing cells with DAPI (blue). (G) Merged channels of single confocal sections of P10 and P14 Control and *Cnot3*-cKO testis showing STRA8 (green) and DDX4 (red) expression with DAPI (blue). (H) Merged channels of single confocal sections of P6, P10 and P14 Control and *Cnot3*-cKO testis showing PCNA (green) and DDX4 (red) expression with DAPI (blue). (I) Merged channels of single confocal sections of P10 and P14 Control and *Cnot3*-cKO testis showing c-CASP3 (green) and DDX4 (red) expression with DAPI (blue).

To better understand the impact of *Cnot3* deletion on SSCs, we analyzed the presence of ZBTB16+ undifferentiated spermatogonia in control and *Cnot3*-cKO mouse testis. The numbers of ZBTB16+ spermatogonia in P6 *Cnot3*-cKO testis were similar to those in control testes (Fig 2C). However, ZBTB16+ spermatogonia were dramatically reduced at P10 and were lost entirely in P14 and P21 in *Cnot3*-cKO mouse testes (Fig 2C), indicating that *Cnot3* deletion led to eventual loss of the undifferentiated spermatogonial pool which includes the foundational adult SSCs. Consistently, the RA-responsive germ cell-specific protein STRA8 (activated in response to RA at the initiation of spermatogonial differentiation) was detected in control but not *Cnot3*-cKO mouse testes at both P10 and P14 (Fig 2G), indicating an absence of differentiating spermatogonia in the *Cnot3*-cKO due to the loss of undifferentiated spermatogonia. Importantly, in *Cnot3*-cKO testes, remaining DDX4+ germ cells were still positive for the proliferation marker PCNA at P6, P10, and P14 (Fig 2H), suggesting that the loss of undifferentiated spermatogonia was not merely caused by defects in germ cell proliferation. Finally, we did not observe increased cleaved (c)-CASP3+ cells (indicative of apoptosis) in *Cnot3*-cKO testes at either P10 or P14 (Fig 2I). Thus, the germ cell loss in cKO testes does not appear to be mediated by apoptosis. Together, these results strongly suggest an essential role for CNOT3 in the maintenance of the nascent spermatogonial populations in the developing testis.

### *Cnot3* deletion impaired the maintenance of steady-state spermatogenesis

As described above, the spermatogonial pool in the developing testis is heterogenous and comprised of SSCs, undifferentiated progenitor, and differentiating spermatogonia. Therefore, we performed scRNA-seq to examine the impact of *Cnot3* deletion on these distinct spermatogonial subpopulations. We used antibodies against surface markers CD9 and THY1 in a fluorescence-activated cell sorting (FACS) approach to enrich for germ cells from freshly prepared P8 control and *Cnot3*-cKO testis single cell suspensions. We performed scRNA-seq using the 10X genomics platform on 4027 cells and 4212 cells from control and *Cnot3*-cKO testes, respectively.

To identify the germ cell populations in these samples, we combined our data with published datasets from P3, P6, P10, and P15 control testes (Ernst et al., 2019; Grive et al., 2019; Law *et al*., 2019). In the combined dataset, we first identified all the germ cells based on *Dazl and Ddx4* (germ cell markers) (Suppl Fig 3A-B). We then used unsupervised clustering and Uniform Manifold Approximation and Projection (UMAP) plots to separate the germ cells into 13 distinct cell populations (Suppl Fig 4A-B). Based on marker gene expression and the developmental time, we separated germ cell clusters into four groups: SSCs, progenitor spermatogonia, differentiating spermatogonia, and preleptotene spermatocytes (Fig 3 and Suppl Fig 4C). Clusters 0 and 2 first appeared at P3 and become less abundant thereafter (Suppl Fig A-D). Germ cells in these clusters also contained high mRNA levels for *Lhx1*, *Gfra1*, and *Zbtb16*, which are enriched in SSCs (Grisanti et al., 2009; Niedenberger et al., 2015; Oatley et al., 2006) (Suppl Fig 4C). It is noteworthy that we previously reported that *Zbtb16* is expressed in nearly all spermatogonia (including undifferentiated and differentiating), during the first wave of spermatogenesis (Suppl Fig 4C) (Niedenberger *et al*., 2015). Based on these results, these clusters were designated as SSCs (Suppl 4A-C). Clusters 1 and 3 became apparent at P6, and included established undifferentiated progenitor markers *Neurog3* (also termed *Ngn3*), *Ddit4* and *Sox3* (Suppl Fig 4A-C), and thus were classified as undifferentiated progenitor spermatogonia (Hobbs *et al*., 2010; Raverot et al., 2005; Yoshida et al., 2004). Clusters 1, 5, and 10 also appeared at P6 and were more abundant at P15. These clusters were enriched for differentiating spermatogonia markers *Stra8* and *Kit* (Suppl Fig 4A-C) and were thus classified as differentiating spermatogonia (Koubova *et al*., 2006; Schrans-Stassen et al., 1999). Clusters 6, 7, 8, 9, and 11 appeared at P15 and included the meiotic marker *Meioc* (Suppl Fig 5C), and were thus classified as spermatocytes (Abby et al., 2016) (Soh et al., 2017).

**Figure 3.**
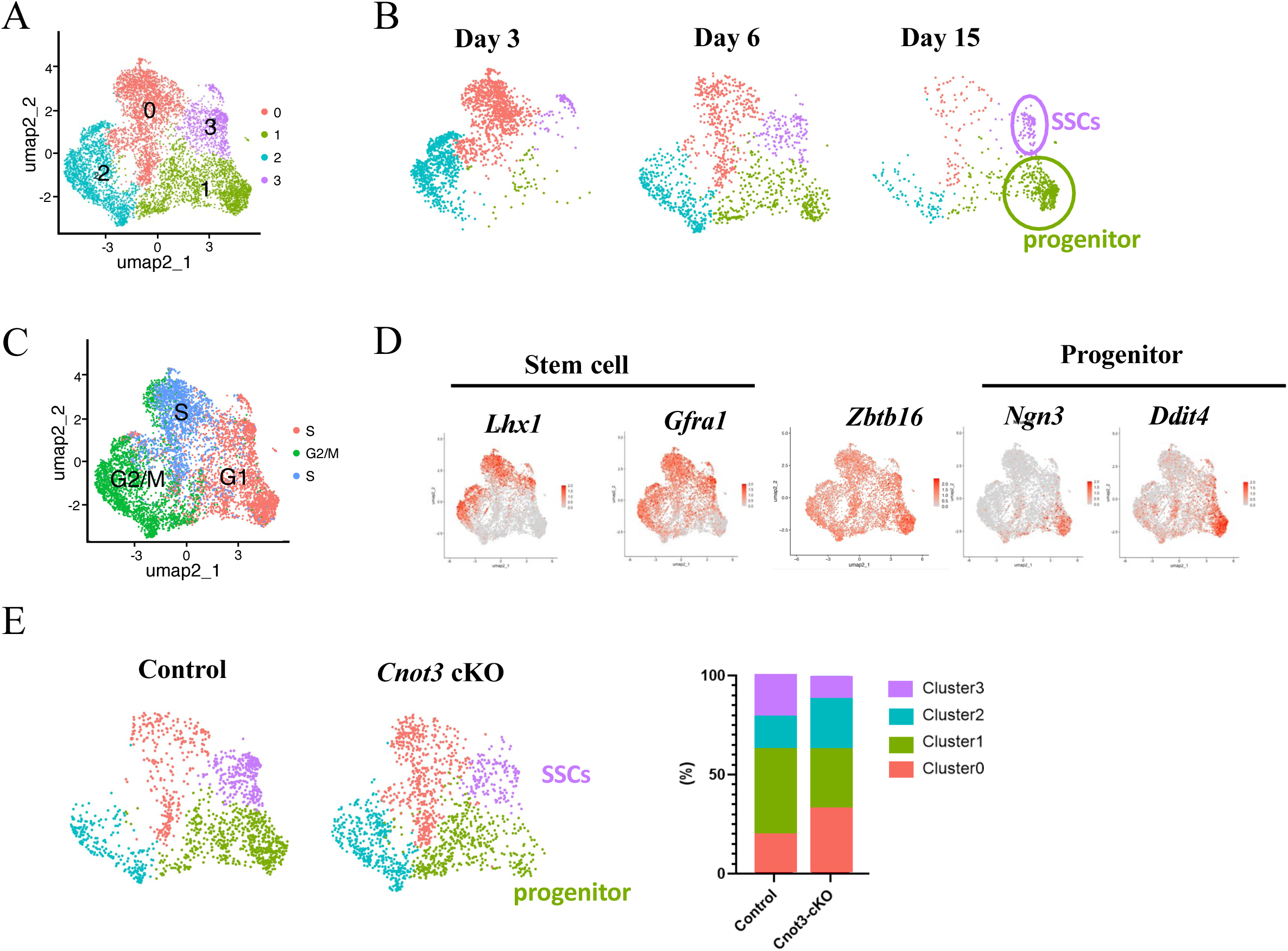
*Cnot3* deletion impairs the maintenance of steady-state spermatogenesis. (A) Reclustering of the group of cells containing SSCs and progenitor cells from the combinaison of our data with published datasets from P3, P6, P10, and P15 stages (Law et al., 2019; Grive et al., 2019, Ernst et al, 2019). (B) Reclustering of the group of cells containing SSCs and progenitor cells from published datasets from P3, P6, and P15 stages, respectively (Law et al., 2019; Grive et al., 2019, Ernst et al, 2019). (C) Highlights of cell cycle phases in reclustered group of cells containing SSCs and progenitor cells from the combinaison of our data with published datasets from P3, P6, P10, and P15 stages (Law et al., 2019; Grive et al., 2019, Ernst et al, 2019). (D) Highlights of Stem cells (*Lhx1* and *Gfra1*) and Progenitor cells (*Ngn3* and *Ddit4*) markers in reclustered group of cells containing SSCs and progenitor cells from the combinaison of our data with published datasets from P3, P6, P10, and P15 stages (Law et al., 2019; Grive et al., 2019, Ernst et al, 2019). (E) Reclustering and quantification of the group of cells containing SSCs and progenitor cells from control and *Cnot3*-cKO.

Since CNOT3 likely plays a critical role in spermatogonia, we focused on germ cell clusters 0, 2, 3, and 4 that contained both SSCs and undifferentiated progenitors. We reanalyzed these cells and generated a new UMAP to separate cells into four new clusters (Fig 3A). In the new UMAP, cells from clusters 0 and 2 were predominantly found in P3 testis samples and gradually disappeared along the development toward P15 (Fig 3A and 3B), suggesting these are spermatogonia undergoing the first wave of spermatogenesis. In addition, the cell cycle status of these cells is in S/G2/M (Fig 3C and Suppl Fig S4D), which is concordant with previous reports that mouse prospermatogonia re-enter the cell cycle before P3 (Drumond *et al*., 2011; Nagano *et al*., 2000). Cell populations in clusters 1 and 3 gradually increased during development into P15 (Fig 3B). Cluster 3 cells contained mRNAs encoding SSC markers LHX1 and GFRA1, while cluster 1 cells contained mRNAs encoding undifferentiated progenitor cell markers NEUROG3 and DDIT4 (Fig 3B and D). Thus, cluster 3 may represent newly formed SSCs required for steady-state spermatogenesis, and cluster 1 represents undifferentiated progenitor spermatogonia.

Comparing the UMAP plots of control and *Cnot3*-cKO germ cells, we found that *Cnot3* deletion significantly decreased cluster 3 cells (=presumed SSCs) and, to a lesser extent, cluster 1 (=presumed undifferentiated progenitor spermatogonia) (Fig 3E). This result is consistent with our conclusions based on immunofluorescence staining (Fig. 2) and provided additional support to the notion that CNOT3 is required for the maintenance of SSCs.

### Cnot3 is required for SSC maintenance *in vitro*

To further test the role of *Cnot3* in SSCs, we derived *in vitro* germ cell cultures enriched for undifferentiated spermatogonia from Cnot3*^flox/flox^* (control SSCs) and [Cnot3*^flox/flox^*; *Ubc*-creER] (*Cnot*-cKO SSCs) mice using an established protocol (Brinster and Avarbock, 1994; Brinster and Zimmermann, 1994; Kubota and Brinster, 2008). Real-time RT-PCR documented that the cultured SSCs highly expressed undifferentiated germ cell markers *Gfra1* and *Zbtb16* but not the Sertoli cell marker *Sox9* (Suppl Fig. 5A). In addition, the mRNA level of *Cnot3* was significantly higher in SSCs than the adherent somatic cells from mouse testis (Suppl Fig. 5A). The SSC identity of the cultured cells was further confirmed by immunofluorescent staining of pan germ cell marker DDX4 and undifferentiated spermatogonia marker ZBTB16 (Suppl Fig. 5B and C). In response to RA treatment, cultured undifferentiated spermatogonia were also able to transit to differentiating spermatogonia, which express both c-KIT and STRA8. (Suppl Fig. 5D). After Tamoxifen (4-OHT) treatment, CNOT3 protein was lost by 72 hours in *Cnot3*-cKO spermatogonia (Suppl Fig. 5B). The deletion of Cnot3 decreased Cnot1 and Cnot2 protein level without affecting the enzymatic module Cnot7, or Cnot8 (Suppl Fig. 5E). We found that *Cnot3* deletion impaired the proliferation and/or viability of cultured undifferentiated spermatogonia (Suppl Fig. 5F). More importantly, the deletion resulted in the down-regulation of key SSC marker genes *Zbtb16*, indicating that CNOT3 is required to maintain undifferentiated spermatogonia, including SSCs (Suppl Fig. 5G). Finally, the *Cnot3*-cKO spermatogonia could not differentiate and activate the differentiating spermatogonia marker c-KIT in response to RA (Suppl Fig. 5H). Therefore, similar to our *in vivo* findings, these results indicate that CNOT3 is required for the maintenance and function of undifferentiated spermatogonia and SSCs *in vitro*.

### *Cnot3* deletion in SSCs affects its transcriptome profile

To understand how *Cnot3* deletion led to dramatic loss of undifferentiated spermatogonia and SSCs, we examined transcriptome changes between control and *Cnot3*-cKO spermatogonia. Since ID4 is an established marker of SSCs in the developing testis (Chan et al., 2014; Cheng et al., 2020; Hermann et al., 2018; Oatley et al., 2011; Velte *et al*., 2019), we generated *Cnot3*^flox/flox^; *Ddx4*-creER; *Id4-eGfp* mice (Helsel et al., 2017; Zheng *et al*., 2016). We isolated single cell suspensions from P8 testes (since *Cnot3* was effectively deleted by this point and CNOT3 protein was undetectable), and collected ID4-EGFP+ spermatogonia by flow cytometry for bulk RNA-seq. Using fold change >2 and adjusted P<0.01 in the RNA-seq analysis, we identified 798 transcripts that were upregulated and 137 that were downregulated in *Cnot3*-cKO testes as compared to controls (Fig 4A, Suppl Table 1). Such a distinct imbalance between the up-and down-regulated mRNA levels is consistent with the known function of *Cnot3* in promoting mRNA degradation (Morita et al., 2011), and suggests CNOT3 normally represses gene expression in SSCs. Based on gene ontology analysis, we found that these up-regulated genes were significantly enriched for RNA metabolic processes and germ cell development (Fig 4B).

**Figure 4.**
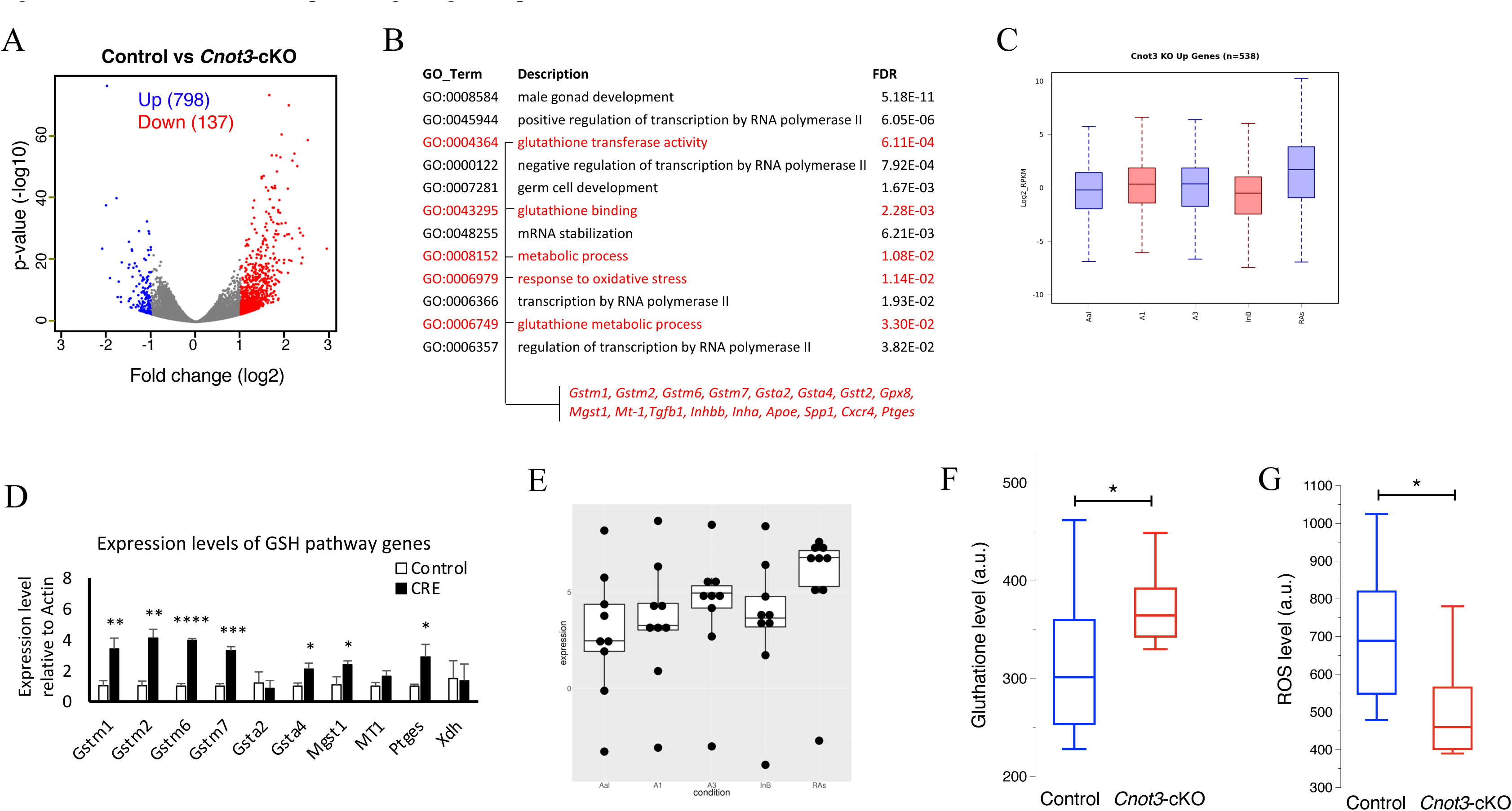
*Cnot3* deletion affects spermatogenic gene expression and GSH/ROS balance. (A) Volcano plot (Log2 Fold change vs -log10 p-value) of RNA-seq data from Control and *Cnot3*-cKO ID4-EGFP+ undifferentiation spermatogonia at P8 (three biological replicates). 798 and 137 genes were upregulated and downregulated, respectively. (B) Gene Ontology analysis of upregulated genes in *Cnot3*-cKO ID4-EGFP+ undifferentiation spermatogonia at P8. Pathway and genes involved in GSH/ROS pathway are indicated in red. (C) Box-plot showing the expression profile of the genes upregulated in *Cnot3*-cKO ID4-EGFP+ SSCs in undifferentiated spermatogonia stages A aligned (Aal) and A1 and differentiating spermatogonia A3, Intermediate B (InB), and preleptotene stages (RAs) (Kirsanov et al., 2023). (D) Expression level of GSH pathway genes relative to expression of *Actin-b* in ID4-EGFP+ spermatogonia from control and cnot3-cKO. Mean ± SEM from three independent measurements are shown (two-tailed unpaired Student’s t test, (^∗^) for p < 0.05, (^∗∗^) for p < 0.01, (^∗∗∗^) for p < 0.001, and (^∗∗∗^) for p < 0.0001). (E) Box-plot showing the expression profile of the GSH pathway genes Gstm1, Gstm2, Gstm6, Gstm7, Gclc, Gsta4, Mgst1, Mt1, Ptges in undifferentiated spermatogonia stages A aligned (Aal) and A1 and differentiating spermatogonia A3, Intermediate B (InB), and preleptotene stages (RAs) (Kirsanov et al., 2023). (F) Whisker and box of Glutathione levels measured by flow cytometry in Control and *Cnot3*-cKO ID4-EGFP+ undifferentiation spermatogonia at P8 (two-tailed unpaired Student’s t test, (^∗^) for p < 0.05). (G) Whisker and box of ROS levels measured by flow cytometry in Control and *Cnot3*-cKO ID4-EGFP+ undifferentiation spermatogonia at P8 (two-tailed unpaired Student’s t test, (^∗^) for p < 0.05).

Next, we collated transcripts increased in *Cnot3*-cKO ID4-EGFP+ spermatogonia and examined their levels during spermatogonial differentiation. This was done using our recently published RNA-seq dataset from germ cells isolated from mice with synchronized steady-state spermatogenesis: undifferentiated spermatogonia and from spermatogonia at the onset (=type A1), midpoint (=type A3), and end (=type intermediate/B) of differentiation, and preleptotene spermatocytes entering meiosis (Kirsanov et al., 2023) (Fig 4C). We found that mRNAs that increased in *Cnot3*-cKO ID4-EGFP+ cells tended to be upregulated between undifferentiated and early/mid-stages (type A1-A3) of spermatogonial differentiation. This observation suggested these *Cnot3*-repressed transcripts are normally upregulated during SSC differentiation. Thus, it is conceivable that CNOT3 normally promotes the degradation of transcripts involved in spermatogonial differentiation to maintain SSC identity and function.

### *Cnot3* deletion upregulates the glutathione redox pathway

Interestingly, among the CNOT3-repressed transcripts in our RNA-seq analyses, we noted that transcripts encoding proteins in the glutathione redox pathway were significantly over-represented (Fig 4B and 4D). The glutathione (GSH) pathway protects cells from oxidative damage by reactive oxygen species (ROS) to promote cell proliferation and differentiation (Meister, 1988) (Sies, 1991). Glutathione S-transferases alpha 4 and m1, (*Gsta4* and *Gstm1)* genes were overexpressed in SSCs compared to differentiating spermatogonia, suggesting a role for the oxidative stress response pathway in regulating the survival and proliferation of SSCs (Kokkinaki et al., 2009; Suzuki et al., 2021). A balance in ROS levels mediated by GSH is critical for spermatogonial maintenance and differentiation – basal ROS levels are critical for self-renewal and proliferation of SSCs (Morimoto et al., 2013; Morimoto et al., 2019), while significantly elevated ROS levels are correlated with male infertility due to their damaging impacts on spermatogonial maintenance and differentiation (Agarwal and Sekhon, 2010; Ishii et al., 2005; Lenzi et al., 1993; Nakamura et al., 2010; Takubo et al., 2006). Thus, we hypothesized *Cnot3* deletion would disrupt the balance between GSH function and ROS levels and thereby impair SSC maintenance. To test the hypothesis, we first examined transcript levels of several genes of the GSH pathway (*Gstm1, Gstm2, Gstm6, Gstm7, Gclc, Gsta4, Mgst1, Mt1, Ptges*) in ID4-EGFP+ cells isolated from control and *Cnot3*-cKO testes. We found that *Cnot3* deletion led to the upregulation of 7 out of the 10 transcripts. Moreover, based on their expression in undifferentiated spermatogonia, A1, A3, and In/B differentiating spermatogonia, and preleptotene spermatocytes from our previous study (Kirsanov *et al*., 2023), these GSH pathway-encoding transcripts were mostly upregulated during spermatogonial differentiation (Fig 4E). These observations suggest CNOT3 represses the GSH pathway in SSCs, and GSH pathway activation may be a necessary step for spermatogonial differentiation. Finally, we examined GSH and ROS levels with flow cytometry using live cell fluorescent dyes in cell suspensions freshly prepared from control and *Cnot3*-cKO P8 testes. In ID4-EGFP-high cells, GSH levels were significantly increased while ROS levels were significantly decreased in the *Cnot3*-cKO samples (Fig 4F-G and Suppl Fig S6), consistent with the role of GSH in preventing ROS accumulation. In summary, our findings revealed that – in the absence of CNOT3 – upregulation of transcripts encoding GSH pathway components led to an accumulation of GSH and a reduction of ROS. This predicts that the reduced ROS level impaired spermatogonial maintenance *in vivo*, resulting in germ cell loss and infertility.

## Discussion

The developmental program of spermatogenesis that is essential for lifelong male fertility relies on the continual function of SSCs, which balance self-renewal divisions with those that produce undifferentiated progenitors that eventually become committed to generating sperm (de Rooij and Russell, 2000; Oatley and Brinster, 2008). The formation and maintenance of these undifferentiated spermatogonial populations depend on a gene expression program regulated by a coordinated balance between both transcriptional and post-transcriptional mechanisms. The germ cell type-specific transcriptome is shaped by the coordinated processes of transcription and RNA decay. Here we discovered that, in mice, CNOT3, a subunit of the CCR4-NOT mRNA deadenylase complex, plays a crucial role in the regulation of SSCs and their potential for self-renewal. CNOT3 is particularly abundant in undifferentiated spermatogonia, and its deletion in spermatogonia led to germ cell loss and infertility. In the absence of CNOT3, transcripts associated with factors involved in SSC differentiation are de-repressed, including those in the glutathione redox pathway which is critical for SSC maintenance. These findings suggest that CNOT3, likely through the CCR4-NOT complex, facilitates the degradation of transcripts linked to differentiation factors essential to maintain the spermatogonial pool and ensure progression of spermatogenesis. This highlights the importance of CCR4-NOT-mediated post-transcriptional gene regulation during male germ cell development.

Poly(A) tails at mRNA 3’ends play an essential role in maintaining mRNA steady-state levels in eukaryotes (Brawerman, 1981; Wickens et al., 1997). Deadenylation, which is the process of removing the poly(A) tail and the initial step in all forms of mRNA decay, is essential for maintaining transcriptome balance (Goldstrohm and Wickens, 2008). The major mRNA deadenylase in eukaryotes is the CCR4-NOT multi-subunit complex that includes the scaffold protein CNOT1 along with the catalytic subunits CNOT8 and CNOT6 (Collart and Panasenko, 2012; 2017). Using genetic approaches, several studies uncovered important roles for CCR4-NOT-mediated mRNA decay during both embryonic and germ cell development. The regulatory CNOT4 subunits are required for post-implantation development and male meiosis (Dai et al., 2021). In whole-body *Cnot6/6l* knockout mice, males were fertile while females were sterile due to defective maternal mRNA degradation during oocyte maturation (Berthet et al., 2004; Sha et al., 2018). Deletion of *Cnot7* impaired spermatogenesis and male fertility, yet oogenesis progressed normally (Berthet *et al*., 2004). CNOT1, CNOT2, and CNOT3 play crucial roles in maintaining both mouse and human ESCs and prevent differentiation into extraembryonic tissues (Zheng *et al*., 2012). It has been observed that the absence of CNOT3 in mice leads to embryonic lethality during the blastocyst stage, primarily due to the loss of inner cell mass (Morita *et al*., 2011; Neely *et al*., 2010). The CNOT proteins can play different roles depending on the cell type context. This suggests that the CCR4-NOT complex subunits target the regulation of different subsets of transcripts depending on the cellular environment. Our results position CNOT3 as a key player in promoting SSC self-renewal. This is achieved, at least in part, through a balanced production of ROS through the regulation of GSH activity.

There are several examples of mRNA binding proteins that interact with the CCR4-NOT deadenylase complex in SSCs to promote the degradation of mRNAs involved in spermatogonial differentiation. This activity is proposed to prevent premature differentiation and ensure that they remain self-renewing. NANOS2, an RNA binding protein that promotes RNA degradation, is only expressed in SSCs in the postnatal testis and mice deficient in NANOS2 have a progressive loss of SSCs leading to germline extinction (Sada et al., 2009) (Codino et al., 2021; Suzuki et al., 2010). In mice, *Dnd1* is expressed from the origin of the germline in primordial germ cells (PGCs) through postnatal spermatogonia. DND1 directly recruits the CCR4-NOT complex to destabilize mRNAs encoding factors involved in the positive regulation of apoptosis, pluripotency, and inflammation. DND1 function is required for the survival of PGCs and SSCs and, in some genetic backgrounds, to prevent the development of testicular germ cell tumors (Cook et al., 2009; Cook et al., 2011; Sakurai et al., 1995; Suzuki et al., 2016; Yamaji et al., 2017).

Our transcriptome analyses showed that transcripts encoding factors targeted by CNOT3 included those in the glutathione redox pathway. Elevated ROS levels can cause oxidative stress and negatively affect male fertility by impacting SSC survival and function (Ishii *et al*., 2005; Nakamura *et al*., 2010; Takubo *et al*., 2006). However, basal levels of ROS are also required for SSC self-renewal (Morimoto *et al*., 2013). Moreover, suppression of ROS levels showed reduced SSC proliferation, while hydrogen peroxide treatment increased SSC self-renewal (Morimoto *et al*., 2013; Morimoto *et al*., 2019). Here, we observed that in SSCs, CNOT3 prevents the elevation of glutathione levels and the decrease of ROS levels, thus protecting the integrity of SSC function. This present study on the postnatal male germline taken together with our previous work in ESCs (Hu *et al*., 2009; Zheng *et al*., 2012) suggest that by targeting distinct transcript subsets, CNOT3 functions at post-transcriptional levels to protect the identity and self-renewal potential of stem cells.

The CCR4-NOT complex plays a multifunctional role at both the transcriptional and post-transcriptional levels and is involved in complex developmental programs. The function of the CCR4-NOT complex is influenced by its interaction with various factors, including mRNA binding proteins, BTG/Tob family factors, and miRNA silencing pathways. In the future, it will be important to understand how the CCR4-NOT complex determines specific target mRNA subsets for degradation.

## Experimental procedures

### Animals and Tamoxifen treatment

*Cnot3*^flox/flox^ and *Ddx4*-CreER transgenic mice were bred to generate *Cnot3*^flox/flox^; *Ddx4*-creER mice. *Cnot3*^flox/flox^; *Ddx4*-creER, *Id4-eGfp* mice were derived by breeding *Cnot3*^flox/flox^; *Ddx4*-creER and *Id4-eGfp* mice (Zheng *et al*., 2016). The genotypes of offspring were determined by qRT-PCR by Transnetyx, Inc. (Zheng *et al*., 2016). For adult mice, 4-week-old *Cnot3*^flox/flox^; *Ddx4*-creER mice were dosed with 200 mg/kg tamoxifen for three consecutive days. Testes were collected from both 6- and 8-week-old mice. Neonatal mice were given daily subcutaneous injections (25 ga needles) of 25 μg of tamoxifen from P1-3. Testes were collected at P6, P8, P10, P14, and P21. All animal procedures were approved by the National Institutes of Health Animals Care and Use Committee and were performed in accordance with an approved National Institute of Environmental Health Sciences animal study proposal.

### Breeding study

Adult males (8-weeks) were housed individually, and two 6-week-old female mice (C57B1/6) were placed with each male for 1 month. After each of the 1-month mating periods, females were held for an additional three weeks to quantify the number of litters and offspring produced.

### *In vitro* SSC culture

SSCs were derived from *Cnot3^flox/flox^* and *Cnot3^flox/flox^; Ddx4-creER* mice that were bred to a DBA/2 background for more than 4 generations following the published protocol (Kubota and Brinster, 2008). Briefly, testes were dissected from mice at the age of P5-P8. After tunica was removed, seminiferous tubules were digested with 7 mg/ml DNase I solution and 0.25% Trypsin-EDTA into a single-cell suspension. Then, SSCs were enriched by 30% Percoll fractionation and then plated on mitomycin C-treated STO cells (SNL76/7 cells) (McMahon and Bradley, 1990). SSCs were maintained in MEMα-based Mouse Serum-Free Medium (SFM) (Kubota and Brinster, 2008). To induce Cnot3 deletion, we added 4-hydroxytamoxifen freshly prepared in ethanol as per manufacturer instructions (4-OHT) (Stem Cell Technologies, 72662) to the medium at 0.1µM at the indicated time points.

The survival curve of control and *Cnot3*-cKO SSCs was determined as follow; cells were treated with vehicle or 4-OHT for 48 hours. Then cells were replated at 2.5 x 10^5^ cells/ well. Cell numbers were determined using a hemocytometer at the indicated timepoints.

To induce SSCs to enter meiosis, cells were treated with 4-OHT or vehicle for 72 hours and with RA (1uM) freshly prepared in DMSO (Cat# R2625, Sigma-Aldrich) for 24 hours before immunofluorescence staining on day 5.

### Tissue processing

Tissues were fixed with 4% paraformaldehyde in 1X PBS overnight at 4°C. For frozen sections, tissues were dehydrated with 30% sucrose and incubated at 4°C until tissue sinks, embedded in O.C.T., and cryosectioned at 10 μm onto glass microscope slides that were then stored at -80°C until use. For paraffin sections, tissues were dehydrated through an ethanol gradient and embedded in paraffin wax using standard methods. Paraffin-embedded specimens were sectioned at 5 μm onto glass microscope slides that were then stored at room temperature until use.

### Histology

5 μm paraffin sections were stained with hematoxylin and eosin (H&E) using standard methods by the Immunohistochemistry Support-Pathology support Group at NIEHS. H&E-stained slides were scanned using Leica Biosystems digital slide scanner by the Imaging Sciences and Artificial Intelligence Group at NIEHS. Representative images were captured using ImageScope through eSlide Manager (Leica Biosystems).

### Immunohistochemistry (IHC)

Paraffin sections were incubated with antigen unmasking solution (H-3300, Vector Laboratories) for antigen retrieval and 3% H_2_O_2_ (H325, Fisher Scientific) to inactivate endogenous peroxidase. Sections were incubated with the blocking reagent (goat serum) and followed with homemade Rabbit anti-mouse CNOT3 (1:1,500) at room temperature for 1 hour (Zheng *et al*., 2012). Sections were washed three times and then incubated with Biotinylated Goat Anti-Rabbit IgG (H+L) for 30 min at room temperature. Then, ABComplex/HRP (VECTASTAIN ABC HRP Kit, Vector Laboratories) and DAB substrate (SK4100, Vector Laboratories) were applied to the sections at room temperature. The sections were counterstained with hematoxylin (HHS16, Sigma), dehydrated, and mounted for imaging. Slides were scanned using Leica Biosystems digital slide scanner by Imaging Sciences and Artificial Intelligence Group at NIEHS. Representative images were captured using ImageScope through eSlide Manager (Leica Biosystems).

### Immunofluorescence (IF) staining

Cryosections were hydrated with 1X PBS and then permeabilized in blocking buffer (1X PBS + 3% BSA + 0.1% Triton X-100) for 30 min at room temperature. Antibodies were applied at appropriate dilutions in the blocking buffer and incubated overnight at 4°C. The slides were washed the next day (1X PBS, 0.05% Triton X-100) for 5 min, followed by two 10 min washes in 1X PBS. Primary antibodies were omitted in negative controls. The primary antibodies were detected by incubating slides for 2 hr at room temperature with corresponding secondary antibodies. After incubation, the slides were washed in (1X PBS, 0.05% Triton X-100) for 5 min, followed by two 10 min washes in 1X PBS, and counterstained with DAPI. Vectashield (Cat#H-1000-10, Vector Laboratories) was used as antifading mounting solution. For paraffin sections, sections were deparaffinized and rehydrated prior to antigen retrieval and standard immunostaining procedures. To perform IF staining on SSCs, SSCs were seeded on STO feeders which were prepared 1 day before on Millicell EZ SLIDE (PEZGS0816, Millipore). SSCs were treated with 4-OHT at 0.1µM or RA at 1 µM or vehicle for the indicated time. Cells were fixed with 4% paraformaldehyde in PBS for 10 min at room temperatures prior to standard immunostaining procedures. Confocal images were taken with a Zeiss LSM 710 inverted microscope (Carl Zeiss Inc, Oberkochen, Germany) using the 405nm, 488nm, and 594nm laser lines for excitation. A Plan-APOCHROMAT 63X/1.4 Oil DIC or a Plan-APOCHROMAT 20x/0.8 objective was used for image collection.

## Antibodies

| Target | Application | Vendor | Host |
| --- | --- | --- | --- |
| CNOT3 | IF, WB | Proteintech (11135-1-AP) | Rabbit |
| CNOT3 | IHC | Zheng et al., 2012 | Rabbit |
| CNOT1 | WB | Proteintech (14276-1-AP) | Rabbit |
| CNOT2 | WB | Proteintech (10313-1-AP) | Rabbit |
| CNOT7 | WB | Cell signaling (86665S) | Rabbit |
| CNOT8 | WB | Invitrogen (PA5-13451) | Rabbit |
| β-Actin | WB | Sigma (A5441) |  |
| ZBTB16 | IF | R&D (AF2944) | Goat |
| DDX4 | IF | R&D (AF2030) | Goat |
| PCNA | IF | Santa Cruz (sc-7907) | Rabbit |
| c-CASP3 | IF | CST(9661S) | Rabbit |
| SOX9 | IF | Millipore(AB5535) | Rabbit |
| STRA8 | IF | Abcam (ab49602) | Rabbit |
| TRA98 | IF | Abcam (ab82527) | rat |
| c-KIT | IF | CST (3074) | Rabbit |
| anti-Rabbit IgG, Alexa Fluor™-488 | IF | ThermoFisher (A-21206) | Donkey |
| anti-Goat IgG, Alexa Fluor™-594 | IF | ThermoFisher (A-11058) | Donkey |
| anti-rat IgG, Alexa Fluor™-594 | IF | ThermoFisher (A-21209) | Donkey |
| ECL anti-Rabbit IgG, HRP | WB | Amersham (NA934-V) | Donkey |

### Immunoblotting

*In vitro* cultured SSCs were lysed in a NuPAGE LDS Sample Buffer (Invitrogen, NP0007) at the same cell concentration. An equal amount of proportion was loaded in each lane.

### FACS isolation of ID4-EGFP cells

Testes were dissected from Tamoxifen-treated neonatal mice (*Cnot3*^flox/flox^;*Id4-eGfp*, and *Cnot3*^flox/flox^; *Ddx4*-creER, ID4-EGFP). Tunica were removed and released seminiferous tubules were placed in a dish with 4.5 ml 0.25% Trypsin (25200056, ThermoFisher) and 0.5 ml Accutase (A1110501, ThermoFisher). Tubules were dissociated by pipetting up and down several times and incubated in a 37°C CO_2_ incubator for 5 min. Another 1 ml Accutase was added, and pipetting and incubation steps were repeated twice until the clumps were dispersed into a single-cell suspension. The single cell suspension was placed on ice, and ID4-EGFP+ cells were collected by a BD FACSAria II Cell Sorter (BD Biosciences). ID4-EGFP+ cell pellets were centrifuged at 300g for 20 min and stored in RNA lysis buffer (Cat# 7326820, Bio-Rad, Aurum™ Total RNA Mini Kit) at -80°C until use.

### The cellular level of ROS and Glutathione measurement

Single-cell suspensions were prepared from P8 ID4-EGFP-control and ID4-EGFP-*Cnot3*-cKO testes. The cells were stained with CellROX™ Deep Red Flow Cytometry Assay Kit (C10491, ThermoFisher) for ROS level and stained with ThiolTracker™ Violet (glutathione detection reagent) (T10095, ThermoFisher) for reduced glutathione according to the manufacturer’s protocol. The cells were analyzed on a BD LSRFortessa™ Cell Analyzer (BD Biosciences). All measurements were independently performed and at least in triplicate. Differences between samples were assessed with the two-tailed unpaired Student’s t-test (QuickCalcs, http://www.graphpad.com/quickcalcs/index.cfm). Statistical significance was indicated as (^∗^) for p < 0.05, (^∗∗^) for p < 0.01, (^∗∗∗^) for p < 0.001, and (ns) for p > 0.05.

### Real time RT-PCR

Total RNA was isolated with TRIZOL reagent (Invitrogen) and treated with TURBO DNA-free (Ambion) as per manufacturer’s recommendations. cDNA synthesis was produced with oligo dT primers and SuperScript III Reverse Transcriptase (Invitrogen). The cDNAs were diluted in water for downstream PCR reactions using appropriate primers and SsoFast EvaGreen Supermix (Bio-Rad) and performed in triplicate. Real-time PCR Bio-Rad CFx96 Real-Time PCR System was used to detect Eva Green. Standard curves were generated using a serial dilution, and actin was used for normalization. Relative quantitation was analyzed using ΔΔCt methods. All measurements were independently performed and at least in triplicate. Differences between samples was assessed with the two-tailed unpaired Student’s t-test (QuickCalcs, http://www.graphpad.com/quickcalcs/index.cfm). Statistical significance was indicated as (^∗^) for p < 0.05, (^∗∗^) for p < 0.01, (^∗∗∗^) for p < 0.001, and (ns) for p > 0.05.

### Real time RT-PCR primers

*Cnot1-F* GACTCGCTCTCGCTGGCCTTG

*Cnot1-R* GCCTGTCTGCCTCAGGACCGTG

*Cnot2-F* CCAACAGAAGCTCGCCAAGC

*Cnot2-R* CCTGATTCCTGTTCATTCCAAATCCAG

*Cnot3-F* AGAGGCCGATCTACAGATAGTGA

*Cnot3-R* GACAGGCTTGGAGCCATTT

*Gfra1-F* CACTCCTGGATTTGCTGATGT

*Gfra1-R* AGTGTGCGGTACTTGGTGC

*Zbtb16-F* GACGCACTACAGGGTTCACA

*Zbtb16-R* GCTTGATCATGGCCGAGTAG

*Sox9-F* ATCTGCACAACGCGGAGCTCA

*Sox9-R* CTCTTCTCGCTCTCGTTCAGCAG

*Gstm1-F* GCAGCTCATCATGCTCTGTT

*Gstm1-R* TTTTCTCAGGGATGGTCTTCA

*Gstm2-F* AGTTGGCCATGGTTTGCTAC

*Gstm2-R* AGCTTCATCTTCTCAGGGAGAC

*Gstm6-F* ATGGGCATGCTTTGCTACA

*Gstm6-R* GGAACTCCGAGTAGAGTTTCAGC

*Gstm7-F* GAGAGGAACCAAGTGTTTGAGGC

*Gstm7-R* TTGGGAGGAAGCGACTGGTCTT

*Gsta2-F* GAGCTTGATGCCAGCCTTCTGA

*Gsta2-R* TTCTCTGGCTGCCAGGATGTAG

*Gsta4-F* GATGATTGCCGTGGCTCCATTTA

*Gsta4-R* CTGGTTGCCAACGAGAAAAGCC

*MGST1-F* GCCAATCCAGAAGACTGTGTAGC

*MGST1-R* AGGAGGCCAATTCCAAGAAATGG

*MT1-F* ACCTCCTTGCAAGAAGAGCTGCT

*MT1-R* GCTGGGTTGGTCCGATACTAT T

*PTGES-F* TCCAGTATTACAGGAGTGACCCAG

*PTGES-R* CCGAGGAAGAGGAAAGGATAGATT

*Xdh-F* GCTCTTCGTGAGCACACAGAAC

*Xdh-R* CCACCCATTCTTTTCACTCGGAC

*bACTIN -F* TCCAGCCTTCCTTCTTGGGTAT

*bACTIN -R* TCTTTACGGATGTCAACGTCACA

### RNA-Seq data Analysis

Reads were quality-filtered, only keeping those with a mean Phred quality score of 20 or greater. Reads were aligned to the mm10 assembly using STAR version 2.6.0c (Dobin et al., 2013). Gene read counts were obtained using the GENCODE annotation version M24 and the featureCounts tool from the Subread package version 1.5.1 (Liao et al., 2014). Differentially expressed genes were identified using DESeq2 version 1.14.1 (Love et al., 2014). GO enrichment analysis was performed using goSTAG (Bennett and Bushel, 2017) version 1.12.1.

### Single-cell RNA seq data Analysis

1. **scRNA-seq data processing**. Raw read processing was carried out using the Cell Ranger Single-Cell Software Suite (version 3.0.1, 10X Genomics Inc., CA, USA). Briefly, the demultiplexed FASTQ files (paired-end, Read 1: 150bp, Read 2:150bp) were generated using the CellRanger mkfastq command. The primary data analyses which included alignment, filtering, barcode counting and UMI quantification for determining gene transcript counts per cell (generated a gene-barcode matrix), quality control, clustering and statistical analysis were performed using CellRanger count command. Gene were annotated using Ensembl build 93.
2. **Single-cell gene expression quantification and filtering**. Raw gene expression matrices generated per sample using CellRanger were imported into R (version 4.0.0) and converted to a Seurat object using the Seurat R package (version 3.4) (Satija et al., 2015). Dead cells and doublets were removed. The first, the total number of UMIs and genes, and percentage of UMIs derived from mitochondrial genome for each cell were counted. Then, Cells which had over 10% UMIs derived from mitochondrial genome were discarded. Next, the upper bound was calculated as mean plus two standard deviation (SD) and the lower bound as mean minus two SD for both the total UMIs and genes, respectively. Finally, Cells with total UMIs or genes outside of the upper and lower bounds were removed.
3. **Data integration (CCA) and determination of the major cell types**. The remaining cells were integrated together, and batch effects were correct using CCA of Seurat (Satija *et al*., 2015). The main cell types were identified on the basis of predicted and known marker genes acquired from the SingleR (https://github.com/LTLA/SingleR) and CellMarker database (http://biocc.hrbmu.edu.cn/CellMarker/).
4. **Data integration of germ cell and determination of the major cell types**. All germ cells raw expression profiles were extracted and integrated together and batch effects were correct using RunFastMNN from R package-“batchelor” (Haghverdi et al., 2018). Firstly, for each sample, gene expression matrices were normalized to total cellular read count and Cell-Cycle scores were calculated using Seurat CellCycleScoring function. Then, Seurat SCTransform function was applied for the normalized data to remove cell cycle effect and select 2500 highly variably genes (HVG) for downstream analysis. We checked HVG and removed mitochondria genes. Following that, We scaled and re-calculated PCA for cleaned HVG. The RunUMAP function was then applied to do the Uniform Manifold Approximation and Projection (UMAP) dimensional reduction. The FindNeighbors constructed a Shared Nearest Neighbor (SNN) Graph, and FindClusters function with “resolution = 0.5” parameter was carried out to cluster cells into different groups. The main cell types were identified on the basis of predicted and known marker genes.

For SSC data analysis, all SSC raw expression profiles were extracted and integrated together and batch effects were correct using RunFastMNN. following analysis is same as for all germ cells.

1. **Identification of marker genes and differential expression genes (DEG).** To identify marker genes for these cell types, we compared the gene expression values of cells from the cluster of interest to that of cells from the rest of clusters using the Seurat FindMarkers function with default parameter of “MAST” (Finak et al., 2015) test. Marker genes were defined based on the following criteria: 1) the average expression value in the cluster of interest was at least 1.2-fold higher than the average expression in the rest of clusters; 2) there are greater than 10% of cells in the cluster of interest which were detectable; and 3) marker genes should have the highest mean expression in the cluster of interest compared to the rest of clusters.

To calculate DEG between two group of cells e.g. N-AML vs healthy donors, Seurat FindMarkers function with method “MAST” were applied for two group of cells with parameter “min.pct = 0.01, logfc.threshold = 0.01”.

For marker genes and DEG lists, GO and pathway analyses were performed by.

1. **Gene set enrichment analysis.** For marker genes and DEG lists, GO and pathway analyses were performed by R package ClusterProfile (V3.18.1) (Wu et al., 2021). The terms with false discovery rate (FDR) <0.05 were regarded as significant enrichment.

## Acknowledgements

The authors wish to thank the NIEHS Epigenomics and DNA sequencing Core, Flow Cytometry Center, Fluorescence Microscopy and Imaging and, Animal facilities for assistance.

This study was supported in part by the Intramural Research Program of the NIH, National Institute of Environmental Health Sciences Z01ES102745 (to G.H.). C.B.D was the recipient of the NIH grants R01HD09003 and R21HD105963.

## Author contributions

Q.C. and G.H. conceived and designed the experiments. Q.C., S.M. and B.L.L. performed the experiments. B.B. and X.X. performed the informatics analysis. S.M., Q.C. and G.H. wrote the paper. O.K. and C.B.G. contributed expertise and reagents.

## References

1. Abby, E., Tourpin, S., Ribeiro, J., Daniel, K., Messiaen, S., Moison, D., Guerquin, J., Gaillard, J.C., Armengaud, J., Langa, F., et al. (2016). Implementation of meiosis prophase I programme requires a conserved retinoid-independent stabilizer of meiotic transcripts. Nat Commun 7, 10324. 10.1038/ncomms10324.

2. Agarwal, A., and Sekhon, L.H. (2010). The role of antioxidant therapy in the treatment of male infertility. Hum Fertil (Camb) 13, 217–225. 10.3109/14647273.2010.532279.

3. Bennett, B.D., and Bushel, P.R. (2017). goSTAG: gene ontology subtrees to tag and annotate genes within a set. Source Code Biol Med 12, 6. 10.1186/s13029-017-0066-1.

4. Berthet, C., Morera, A.M., Asensio, M.J., Chauvin, M.A., Morel, A.P., Dijoud, F., Magaud, J.P., Durand, P., and Rouault, J.P. (2004). CCR4-associated factor CAF1 is an essential factor for spermatogenesis. Mol Cell Biol 24, 5808–5820. 10.1128/MCB.24.13.5808-5820.2004.

5. Brawerman, G. (1981). The Role of the poly(A) sequence in mammalian messenger RNA. CRC Crit Rev Biochem 10, 1–38. 10.3109/10409238109114634.

6. Brinster, R.L., and Avarbock, M.R. (1994). Germline transmission of donor haplotype following spermatogonial transplantation. Proc Natl Acad Sci U S A 91, 11303–11307. 10.1073/pnas.91.24.11303.

7. Brinster, R.L., and Zimmermann, J.W. (1994). Spermatogenesis following male germ-cell transplantation. Proc Natl Acad Sci U S A 91, 11298–11302. 10.1073/pnas.91.24.11298.

8. Buaas, F.W., Kirsh, A.L., Sharma, M., McLean, D.J., Morris, J.L., Griswold, M.D., de Rooij, D.G., and Braun, R.E. (2004). Plzf is required in adult male germ cells for stem cell self-renewal. Nat Genet 36, 647–652. 10.1038/ng1366.

9. Chan, F., Oatley, M.J., Kaucher, A.V., Yang, Q.E., Bieberich, C.J., Shashikant, C.S., and Oatley, J.M. (2014). Functional and molecular features of the Id4+ germline stem cell population in mouse testes. Genes Dev 28, 1351–1362. 10.1101/gad.240465.114.

10. Cheng, K., Chen, I.C., Cheng, C.E., Mutoji, K., Hale, B.J., Hermann, B.P., Geyer, C.B., Oatley, J.M., and McCarrey, J.R. (2020). Unique Epigenetic Programming Distinguishes Regenerative Spermatogonial Stem Cells in the Developing Mouse Testis. iScience 23, 101596. 10.1016/j.isci.2020.101596.

11. Codino, A., Turowski, T., van de Lagemaat, L.N., Ivanova, I., Tavosanis, A., Much, C., Auchynnikava, T., Vasiliauskaite, L., Morgan, M., Rappsilber, J., et al. (2021). NANOS2 is a sequence-specific mRNA-binding protein that promotes transcript degradation in spermatogonial stem cells. iScience 24, 102762. 10.1016/j.isci.2021.102762.

12. Collart, M.A., and Panasenko, O.O. (2012). The Ccr4--not complex. Gene 492, 42–53. 10.1016/j.gene.2011.09.033.

13. Collart, M.A., and Panasenko, O.O. (2017). The Ccr4-Not Complex: Architecture and Structural Insights. Subcell Biochem 83, 349–379. 10.1007/978-3-319-46503-6_13.

14. Cook, M.S., Coveney, D., Batchvarov, I., Nadeau, J.H., and Capel, B. (2009). BAX-mediated cell death affects early germ cell loss and incidence of testicular teratomas in Dnd1(Ter/Ter) mice. Dev Biol 328, 377–383. 10.1016/j.ydbio.2009.01.041.

15. Cook, M.S., Munger, S.C., Nadeau, J.H., and Capel, B. (2011). Regulation of male germ cell cycle arrest and differentiation by DND1 is modulated by genetic background. Development 138, 23–32. 10.1242/dev.057000.

16. Costoya, J.A., Hobbs, R.M., Barna, M., Cattoretti, G., Manova, K., Sukhwani, M., Orwig, K.E., Wolgemuth, D.J., and Pandolfi, P.P. (2004). Essential role of Plzf in maintenance of spermatogonial stem cells. Nat Genet 36, 653–659. 10.1038/ng1367.

17. Dai, X.X., Jiang, Y., Gu, J.H., Jiang, Z.Y., Wu, Y.W., Yu, C., Yin, H., Zhang, J., Shi, Q.H., Shen, L., et al. (2021). The CNOT4 Subunit of the CCR4-NOT Complex is Involved in mRNA Degradation, Efficient DNA Damage Repair, and XY Chromosome Crossover during Male Germ Cell Meiosis. Adv Sci (Weinh) 8, 2003636. 10.1002/advs.202003636.

18. de Kretser, D.M.L., K L; Meinhardt, A; Simorangkir, D; Wreford, N (1998). Spermatogenesis. Human Reproduction 13 (*1*), 1–8. 0.1093/humrep/13.suppl_1.1.

19. de Rooij, D.G. (2001). Proliferation and differentiation of spermatogonial stem cells. Reproduction 121, 347–354. 10.1530/rep.0.1210347.

20. de Rooij, D.G., and Russell, L.D. (2000). All you wanted to know about spermatogonia but were afraid to ask. J Androl 21, 776–798.

21. Dobin, A., Davis, C.A., Schlesinger, F., Drenkow, J., Zaleski, C., Jha, S., Batut, P., Chaisson, M., and Gingeras, T.R. (2013). STAR: ultrafast universal RNA-seq aligner. Bioinformatics 29, 15–21. 10.1093/bioinformatics/bts635.

22. Drumond, A.L., Meistrich, M.L., and Chiarini-Garcia, H. (2011). Spermatogonial morphology and kinetics during testis development in mice: a high-resolution light microscopy approach. Reproduction 142, 145–155. 10.1530/REP-10-0431.

23. Du, G., Wang, X., Luo, M., Xu, W., Zhou, T., Wang, M., Yu, L., Li, L., Cai, L., Wang, P.J., et al. (2020). mRBPome capture identifies the RNA-binding protein TRIM71, an essential regulator of spermatogonial differentiation. Development 147. 10.1242/dev.184655.

24. Ernst, C., Eling, N., Martinez-Jimenez, C.P., Marioni, J.C., and Odom, D.T. (2019). Staged developmental mapping and X chromosome transcriptional dynamics during mouse spermatogenesis. Nat Commun 10, 1251. 10.1038/s41467-019-09182-1.

25. Finak, G., McDavid, A., Yajima, M., Deng, J., Gersuk, V., Shalek, A.K., Slichter, C.K., Miller, H.W., McElrath, M.J., Prlic, M., et al. (2015). MAST: a flexible statistical framework for assessing transcriptional changes and characterizing heterogeneity in single-cell RNA sequencing data. Genome Biol 16, 278. 10.1186/s13059-015-0844-5.

26. Gewiss, R.L., Schleif, M.C., and Griswold, M.D. (2021). The role of retinoic acid in the commitment to meiosis. Asian J Androl 23, 549–554. 10.4103/aja202156.

27. Goldstrohm, A.C., and Wickens, M. (2008). Multifunctional deadenylase complexes diversify mRNA control. Nat Rev Mol Cell Biol 9, 337–344. 10.1038/nrm2370.

28. Green, C.D., Ma, Q., Manske, G.L., Shami, A.N., Zheng, X., Marini, S., Moritz, L., Sultan, C., Gurczynski, S.J., Moore, B.B., et al. (2018). A Comprehensive Roadmap of Murine Spermatogenesis Defined by Single-Cell RNA-Seq. Dev Cell 46, 651–667 e610. 10.1016/j.devcel.2018.07.025.

29. Grisanti, L., Falciatori, I., Grasso, M., Dovere, L., Fera, S., Muciaccia, B., Fuso, A., Berno, V., Boitani, C., Stefanini, M., and Vicini, E. (2009). Identification of spermatogonial stem cell subsets by morphological analysis and prospective isolation. Stem Cells 27, 3043–3052. 10.1002/stem.206.

30. Griswold, M.D. (2016). Spermatogenesis: The Commitment to Meiosis. Physiol Rev 96, 1–17. 10.1152/physrev.00013.2015.

31. Grive, K.J., Hu, Y., Shu, E., Grimson, A., Elemento, O., Grenier, J.K., and Cohen, P.E. (2019). Dynamic transcriptome profiles within spermatogonial and spermatocyte populations during postnatal testis maturation revealed by single-cell sequencing. PLoS Genet 15, e1007810. 10.1371/journal.pgen.1007810.

32. Haghverdi, L., Lun, A.T.L., Morgan, M.D., and Marioni, J.C. (2018). Batch effects in single-cell RNA-sequencing data are corrected by matching mutual nearest neighbors. Nat Biotechnol 36, 421–427. 10.1038/nbt.4091.

33. Helsel, A.R., Yang, Q.E., Oatley, M.J., Lord, T., Sablitzky, F., and Oatley, J.M. (2017). ID4 levels dictate the stem cell state in mouse spermatogonia. Development 144, 624–634. 10.1242/dev.146928.

34. Hermann, B.P., Cheng, K., Singh, A., Roa-De La Cruz, L., Mutoji, K.N., Chen, I.C., Gildersleeve, H., Lehle, J.D., Mayo, M., Westernstroer, B., et al. (2018). The Mammalian Spermatogenesis Single-Cell Transcriptome, from Spermatogonial Stem Cells to Spermatids. Cell Rep 25, 1650–1667 e1658. 10.1016/j.celrep.2018.10.026.

35. Hermann, B.P., Mutoji, K.N., Velte, E.K., Ko, D., Oatley, J.M., Geyer, C.B., and McCarrey, J.R. (2015). Transcriptional and translational heterogeneity among neonatal mouse spermatogonia. Biol Reprod 92, 54. 10.1095/biolreprod.114.125757.

36. Hobbs, R.M., Seandel, M., Falciatori, I., Rafii, S., and Pandolfi, P.P. (2010). Plzf regulates germline progenitor self-renewal by opposing mTORC1. Cell 142, 468–479. 10.1016/j.cell.2010.06.041.

37. Hu, G., Kim, J., Xu, Q., Leng, Y., Orkin, S.H., and Elledge, S.J. (2009). A genome-wide RNAi screen identifies a new transcriptional module required for self-renewal. Genes Dev 23, 837–848. 10.1101/gad.1769609.

38. Ishii, H., Arai, T., Mori, H., Yamada, H., Endo, N., Makino, K., and Fukuda, K. (2005). Protective effects of intracellular reactive oxygen species generated by 6-formylpterin on tumor necrosis factor-alpha-induced apoptotic cell injury in cultured rat hepatocytes. Life Sci 77, 858–868. 10.1016/j.lfs.2004.11.038.

39. Kirsanov, O., Johnson, T.A., Niedenberger, B.A., Malachowski, T.N., Hale, B.J., Chen, Q., Lackford, B., Wang, J., Singh, A., Schindler, K., et al. (2023). Retinoic acid is dispensable for meiotic initiation but required for spermiogenesis in the mammalian testis. Development 150. 10.1242/dev.201638.

40. Kluin, P.M., and de Rooij, D.G. (1981). A comparison between the morphology and cell kinetics of gonocytes and adult type undifferentiated spermatogonia in the mouse. Int J Androl 4, 475–493. 10.1111/j.1365-2605.1981.tb00732.x.

41. Kokkinaki, M., Lee, T.L., He, Z., Jiang, J., Golestaneh, N., Hofmann, M.C., Chan, W.Y., and Dym, M. (2009). The molecular signature of spermatogonial stem/progenitor cells in the 6-day-old mouse testis. Biol Reprod 80, 707–717. 10.1095/biolreprod.108.073809.

42. Koubova, J., Menke, D.B., Zhou, Q., Capel, B., Griswold, M.D., and Page, D.C. (2006). Retinoic acid regulates sex-specific timing of meiotic initiation in mice. Proc Natl Acad Sci U S A 103, 2474–2479. 10.1073/pnas.0510813103.

43. Kubota, H., and Brinster, R.L. (2008). Culture of rodent spermatogonial stem cells, male germline stem cells of the postnatal animal. Methods Cell Biol 86, 59–84. 10.1016/S0091-679X(08)00004-6.

44. Law, N.C., Oatley, M.J., and Oatley, J.M. (2019). Developmental kinetics and transcriptome dynamics of stem cell specification in the spermatogenic lineage. Nat Commun 10, 2787. 10.1038/s41467-019-10596-0.

45. Legrand, J.M.D., and Hobbs, R.M. (2018). RNA processing in the male germline: Mechanisms and implications for fertility. Semin Cell Dev Biol 79, 80–91. 10.1016/j.semcdb.2017.10.006.

46. Lenzi, A., Culasso, F., Gandini, L., Lombardo, F., and Dondero, F. (1993). Placebo-controlled, double-blind, cross-over trial of glutathione therapy in male infertility. Hum Reprod 8, 1657–1662. 10.1093/oxfordjournals.humrep.a137909.

47. Liao, Y., Smyth, G.K., and Shi, W. (2014). featureCounts: an efficient general purpose program for assigning sequence reads to genomic features. Bioinformatics 30, 923–930. 10.1093/bioinformatics/btt656.

48. Love, M.I., Huber, W., and Anders, S. (2014). Moderated estimation of fold change and dispersion for RNA-seq data with DESeq2. Genome Biol 15, 550. 10.1186/s13059-014-0550-8.

49. McMahon, A.P., and Bradley, A. (1990). The Wnt-1 (int-1) proto-oncogene is required for development of a large region of the mouse brain. Cell 62, 1073–1085. 10.1016/0092-8674(90)90385-r.

50. Meister, A. (1988). Glutathione metabolism and its selective modification. J Biol Chem 263, 17205–17208.

51. Morimoto, H., Iwata, K., Ogonuki, N., Inoue, K., Atsuo, O., Kanatsu-Shinohara, M., Morimoto, T., Yabe-Nishimura, C., and Shinohara, T. (2013). ROS are required for mouse spermatogonial stem cell self-renewal. Cell Stem Cell 12, 774–786. 10.1016/j.stem.2013.04.001.

52. Morimoto, H., Kanastu-Shinohara, M., Ogonuki, N., Kamimura, S., Ogura, A., Yabe-Nishimura, C., Mori, Y., Morimoto, T., Watanabe, S., Otsu, K., et al. (2019). ROS amplification drives mouse spermatogonial stem cell self-renewal. Life Sci Alliance 2. 10.26508/lsa.201900374.

53. Morita, M., Oike, Y., Nagashima, T., Kadomatsu, T., Tabata, M., Suzuki, T., Nakamura, T., Yoshida, N., Okada, M., and Yamamoto, T. (2011). Obesity resistance and increased hepatic expression of catabolism-related mRNAs in Cnot3+/-mice. EMBO J 30, 4678–4691. 10.1038/emboj.2011.320.

54. Nagano, R., Tabata, S., Nakanishi, Y., Ohsako, S., Kurohmaru, M., and Hayashi, Y. (2000). Reproliferation and relocation of mouse male germ cells (gonocytes) during prespermatogenesis. Anat Rec 258, 210–220. 10.1002/(SICI)1097-0185(20000201)258:2210::AID-AR103.0.CO;2-X.

55. Nakamura, N., Mori, C., and Eddy, E.M. (2010). Molecular complex of three testis-specific isozymes associated with the mouse sperm fibrous sheath: hexokinase 1, phosphofructokinase M, and glutathione S-transferase mu class 5. Biol Reprod 82, 504–515. 10.1095/biolreprod.109.080580.

56. Neely, G.G., Kuba, K., Cammarato, A., Isobe, K., Amann, S., Zhang, L., Murata, M., Elmen, L., Gupta, V., Arora, S., et al. (2010). A global in vivo Drosophila RNAi screen identifies NOT3 as a conserved regulator of heart function. Cell 141, 142–153. 10.1016/j.cell.2010.02.023.

57. Niedenberger, B.A., Busada, J.T., and Geyer, C.B. (2015). Marker expression reveals heterogeneity of spermatogonia in the neonatal mouse testis. Reproduction 149, 329–338. 10.1530/REP-14-0653.

58. Niimi, Y., Imai, A., Nishimura, H., Yui, K., Kikuchi, A., Koike, H., Saga, Y., and Suzuki, A. (2019). Essential role of mouse Dead end1 in the maintenance of spermatogonia. Dev Biol 445, 103–112. 10.1016/j.ydbio.2018.11.003.

59. Nikolova, D.B., Martinova, Y.S., Seidensticker, M., and Bellve, A.R. (1997). Leukaemia inhibitory factor stimulates proliferation of prospermatogonial stem cells. Reprod Fertil Dev 9, 717–721. 10.1071/r96116.

60. Oatley, J.M., Avarbock, M.R., Telaranta, A.I., Fearon, D.T., and Brinster, R.L. (2006). Identifying genes important for spermatogonial stem cell self-renewal and survival. Proc Natl Acad Sci U S A 103, 9524–9529. 10.1073/pnas.0603332103.

61. Oatley, J.M., and Brinster, R.L. (2006). Spermatogonial stem cells. Methods Enzymol 419, 259–282. 10.1016/S0076-6879(06)19011-4.

62. Oatley, J.M., and Brinster, R.L. (2008). Regulation of spermatogonial stem cell self-renewal in mammals. Annu Rev Cell Dev Biol 24, 263–286. 10.1146/annurev.cellbio.24.110707.175355.

63. Oatley, M.J., Kaucher, A.V., Racicot, K.E., and Oatley, J.M. (2011). Inhibitor of DNA binding 4 is expressed selectively by single spermatogonia in the male germline and regulates the self-renewal of spermatogonial stem cells in mice. Biol Reprod 85, 347–356. 10.1095/biolreprod.111.091330.

64. Raverot, G., Weiss, J., Park, S.Y., Hurley, L., and Jameson, J.L. (2005). Sox3 expression in undifferentiated spermatogonia is required for the progression of spermatogenesis. Dev Biol 283, 215–225. 10.1016/j.ydbio.2005.04.013.

65. Sada, A., Suzuki, A., Suzuki, H., and Saga, Y. (2009). The RNA-binding protein NANOS2 is required to maintain murine spermatogonial stem cells. Science 325, 1394–1398. 10.1126/science.1172645.

66. Sakurai, T., Iguchi, T., Moriwaki, K., and Noguchi, M. (1995). The ter mutation first causes primordial germ cell deficiency in ter/ter mouse embryos at 8 days of gestation. Dev Growth Differ 37, 293–302. 10.1046/j.1440-169X.1995.t01-2-00007.x.

67. Satija, R., Farrell, J.A., Gennert, D., Schier, A.F., and Regev, A. (2015). Spatial reconstruction of single-cell gene expression data. Nat Biotechnol 33, 495–502. 10.1038/nbt.3192.

68. Schrans-Stassen, B.H., van de Kant, H.J., de Rooij, D.G., and van Pelt, A.M. (1999). Differential expression of c-kit in mouse undifferentiated and differentiating type A spermatogonia. Endocrinology 140, 5894–5900. 10.1210/endo.140.12.7172.

69. Sha, Q.Q., Yu, J.L., Guo, J.X., Dai, X.X., Jiang, J.C., Zhang, Y.L., Yu, C., Ji, S.Y., Jiang, Y., Zhang, S.Y., et al. (2018). CNOT6L couples the selective degradation of maternal transcripts to meiotic cell cycle progression in mouse oocyte. EMBO J 37. 10.15252/embj.201899333.

70. Sies, H. (1991). Role of reactive oxygen species in biological processes. Klin Wochenschr 69, 965–968. 10.1007/BF01645140.

71. Soh, Y.Q.S., Mikedis, M.M., Kojima, M., Godfrey, A.K., de Rooij, D.G., and Page, D.C. (2017). Meioc maintains an extended meiotic prophase I in mice. PLoS Genet 13, e1006704. 10.1371/journal.pgen.1006704.

72. Suzuki, A., Igarashi, K., Aisaki, K., Kanno, J., and Saga, Y. (2010). NANOS2 interacts with the CCR4-NOT deadenylation complex and leads to suppression of specific RNAs. Proc Natl Acad Sci U S A 107, 3594–3599. 10.1073/pnas.0908664107.

73. Suzuki, A., Niimi, Y., Shinmyozu, K., Zhou, Z., Kiso, M., and Saga, Y. (2016). Dead end1 is an essential partner of NANOS2 for selective binding of target RNAs in male germ cell development. EMBO Rep 17, 37–46. 10.15252/embr.201540828.

74. Suzuki, S., McCarrey, J.R., and Hermann, B.P. (2021). An mTORC1-dependent switch orchestrates the transition between mouse spermatogonial stem cells and clones of progenitor spermatogonia. Cell Rep 34, 108752. 10.1016/j.celrep.2021.108752.

75. Takubo, K., Hirao, A., Ohmura, M., Azuma, M., Arai, F., Nagamatsu, G., and Suda, T. (2006). Premeiotic germ cell defect in seminiferous tubules of Atm-null testis. Biochem Biophys Res Commun 351, 993–998. 10.1016/j.bbrc.2006.10.145.

76. Velte, E.K., Niedenberger, B.A., Serra, N.D., Singh, A., Roa-DeLaCruz, L., Hermann, B.P., and Geyer, C.B. (2019). Differential RA responsiveness directs formation of functionally distinct spermatogonial populations at the initiation of spermatogenesis in the mouse. Development 146. 10.1242/dev.173088.

77. Wickens, M., Anderson, P., and Jackson, R.J. (1997). Life and death in the cytoplasm: messages from the 3’ end. Curr Opin Genet Dev 7, 220–232. 10.1016/s0959-437x(97)80132-3.

78. Wu, T., Hu, E., Xu, S., Chen, M., Guo, P., Dai, Z., Feng, T., Zhou, L., Tang, W., Zhan, L., et al. (2021). clusterProfiler 4.0: A universal enrichment tool for interpreting omics data. Innovation (Camb) 2, 100141. 10.1016/j.xinn.2021.100141.

79. Yamaji, M., Jishage, M., Meyer, C., Suryawanshi, H., Der, E., Yamaji, M., Garzia, A., Morozov, P., Manickavel, S., McFarland, H.L., et al. (2017). DND1 maintains germline stem cells via recruitment of the CCR4-NOT complex to target mRNAs. Nature 543, 568–572. 10.1038/nature21690.

80. Yoshida, S., Sukeno, M., Nakagawa, T., Ohbo, K., Nagamatsu, G., Suda, T., and Nabeshima, Y. (2006). The first round of mouse spermatogenesis is a distinctive program that lacks the self-renewing spermatogonia stage. Development 133, 1495–1505. 10.1242/dev.02316.

81. Yoshida, S., Takakura, A., Ohbo, K., Abe, K., Wakabayashi, J., Yamamoto, M., Suda, T., and Nabeshima, Y. (2004). Neurogenin3 delineates the earliest stages of spermatogenesis in the mouse testis. Dev Biol 269, 447–458. 10.1016/j.ydbio.2004.01.036.

82. Zheng, X., Dumitru, R., Lackford, B.L., Freudenberg, J.M., Singh, A.P., Archer, T.K., Jothi, R., and Hu, G. (2012). Cnot1, Cnot2, and Cnot3 maintain mouse and human ESC identity and inhibit extraembryonic differentiation. Stem Cells 30, 910–922. 10.1002/stem.1070.

83. Zheng, X., Yang, P., Lackford, B., Bennett, B.D., Wang, L., Li, H., Wang, Y., Miao, Y., Foley, J.F., Fargo, D.C., et al. (2016). CNOT3-Dependent mRNA Deadenylation Safeguards the Pluripotent State. Stem Cell Reports 7, 897–910. 10.1016/j.stemcr.2016.09.007.

84. Zhou, W., Shao, H., Zhang, D., Dong, J., Cheng, W., Wang, L., Teng, Y., and Yu, Z. (2015). PTEN signaling is required for the maintenance of spermatogonial stem cells in mouse, by regulating the expressions of PLZF and UTF1. Cell Biosci 5, 42. 10.1186/s13578-015-0034-x.

