## Supplemental Legends and figures for "*Cnot3* is required for male germ cell development and spermatogonial stem cell maintenance"

#### Supplemental Figure Legends

**Figure S1. *Cnot3* deletion results in germ cell loss and male infertility**

(A) Analyses of the abundance of *Cnot3* transcripts in Spermatogonia (SPG), Spermatocytes (Scytes), Spermatids (STids) and Elongating spermatids from the single-cell RNA seq data generated by Green et al., 2018.

(B) Sections of testis of 8wo WT male showing CNOT3 (brown) and hematoxylin.

(C) Merged channels of single confocal sections of 8wo testis showing CNOT3 (green) expression in ZBTB16-positive (red) in undifferentiated spermatogonia with DAPI (blue).

**Figure S2. Effect of *Cnot3* deletion on germ cell and Sertoli cells in juvenile testis**

Merged channels of single confocal sections of P6, P10, P14 and P21 Control and *Cnot3*-cKO testis showing SOX9 (green) and DDX4 (red) expression with DAPI (blue).

**Figure S3. Identification of the germ cells population based on clustering and marker gene expression**

(A) UMAP projection of the testicular cells from the combination of our data with published datasets from P3, P6, P10, and P15 stages and indication of the Macrophage, Sertoli, Endothelial and germ cells in testicular (Law et al., 2019; Grive et al., 2019, Ernst et al, 2019).

(B) Highlights of of the Macrophage (Adgre1, Csf1r), Sertoli (Sox9, Amhr2), Endothelial (Lyc6c1) and germ cells (Dazl, Ddx4) in testicular in the UMAP projection of the testicular cells from the combination of our data with published datasets from P3, P6, P10, and P15 stages (Law et al., 2019; Grive et al., 2019, Ernst et al, 2019)

**Figure S4. Identification of the germ cells population based on clustering and marker gene expression**

(A) Clustering of the of cells from the combination of our data with published datasets from P3, P6, and P15 stages (Law et al., 2019; Grive et al., 2019, Ernst et al, 2019).

(B) Clustering of the of cells from from the published datasets from P3, P6, and P15 stages, respectively (Law et al., 2019; Grive et al., 2019, Ernst et al, 2019).

(C) Highlights of Stem cells (*Lhx1, Gfra1 and, Etv5*), Progenitor cells (*Ngn3,Ddit4 and, Sox3*), differentiating (*Stra8 and c-Kit*) and maturing (Meioc) spermatogonia markers in clustered group of cells from the combination of our data with published datasets from P3, P6, P10, and P15 stages (Law et al., 2019; Grive et al., 2019, Ernst et al, 2019).

(D) Highlights of cell cycle phases in clustered group of cells from the combination of our data with published datasets from P3, P6, P10, and P15 stages (Law et al., 2019; Grive et al., 2019, Ernst et al, 2019).

**Figure S5. CNOT3 is required for SSC maintenance and differentiation *in vitro***

(A) Expression level of *Cnot1*, *Cnot2*, *Cnot3*, *Gfra1*, *Zbtb16* and *Sox9* genes relative to β-actin in adherent cells and SSCs derived from control testis. Mean ± SEM from three independent measurements are shown.

(B) Single and merged channels of SSCs derived from Control and *Cnot3*-cKO testis showing CNOT3 (green) and DDX4 (red) expression. Cells were treated with tamoxifen for 72 hours.

(C) Single and merged channels of SSCs derived from Control testis showing CNOT3 (green) and ZBTB16 (red) expression.

(D) Single and merged channels of SSCs derived from Control testis showing c-KIT (green) and DDX4 (red) expression (top 2 panels) and STRA8 (green) and DDX4 (red) expression (bottom 2 panels). Cells were treated with retinoic acid (RA, 1μM) or vehicle for 24 hours.

(E) The protein level of CCR4-NOT subunits was detected in Control and *Cnot3*-cKO SSCs. Cells were treated with tamoxifen for 72 hours and collected at indicated timepoints. Feeder cells (STO) as a control.

(F) The survival curve of control and *Cnot3*-cKO SSCs was determined. Cells were treated with vehicle or 4-OHT for 48 hours. Then cells were replated at 2.5 x $10^5$ cells/ well. Cell numbers were determined using a hemocytometer at the indicated timepoints. Mean ± SEM from three independent measurements are shown.

(G) Relative expression level of *Cnot3* and *Zbtb16* genes in control and *Cnot3*-cKO SSCs treated with tamoxifen (4-OHT) for 48 hours at the indicated timepoints. Mean ± SEM from three independent measurements are shown.

(H) Single and merged channels of SSCs derived from *Cnot3*-cKO testis showing c-KIT (green) and DDX4 (red) expression with DAPI. Cells were treated with 4-OHT or vehicle for 72 hours and with RA (1uM) for 24 hours before IF staining on day 5.

**Figure S6. Effect of *Cnot3* deletion on the glutathione and ROS content in ID4-EGFP+ undifferentiation spermatogonia at P8**

(A) Glutathione content in Control and *Cnot3*-cKO ID4-EGFP+ undifferentiation spermatogonia at P8 analyzed by flow cytometry. An increase in Glutathione content resulted in an increase in the fluorescence intensity.

(B) ROS content in content in Control and *Cnot3*-cKO ID4-EGFP+ undifferentiation spermatogonia at P8 analyzed by flow cytometry. A decrease in ROS content resulted in a decrease in the fluorescence intensity.

**Figure S1. *Cnot3* deletion results in germ cell loss and male infertility**

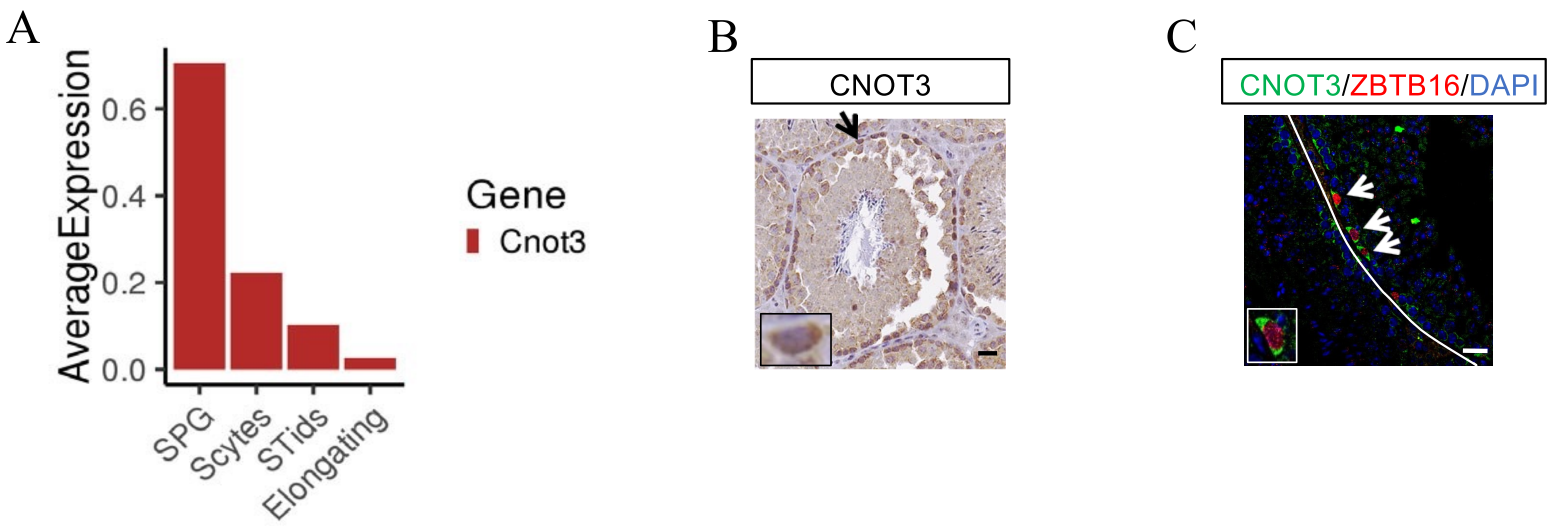

**Figure S2. Effect of *Cnot3* deletion on germ cell and Sertoli cells in juvenile testis**

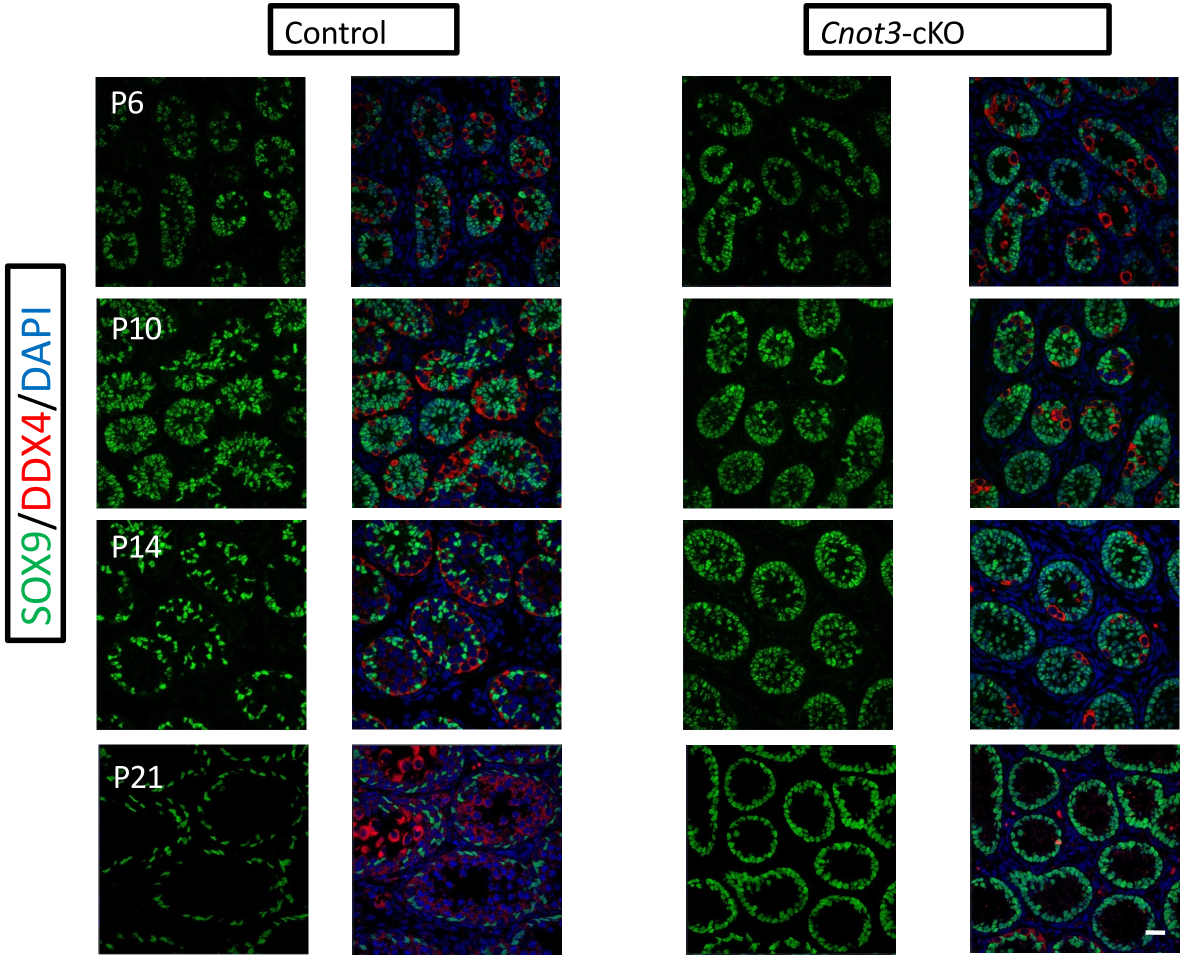

**Figure S3. Identification of the germ cells population based on clustering and marker gene expression**

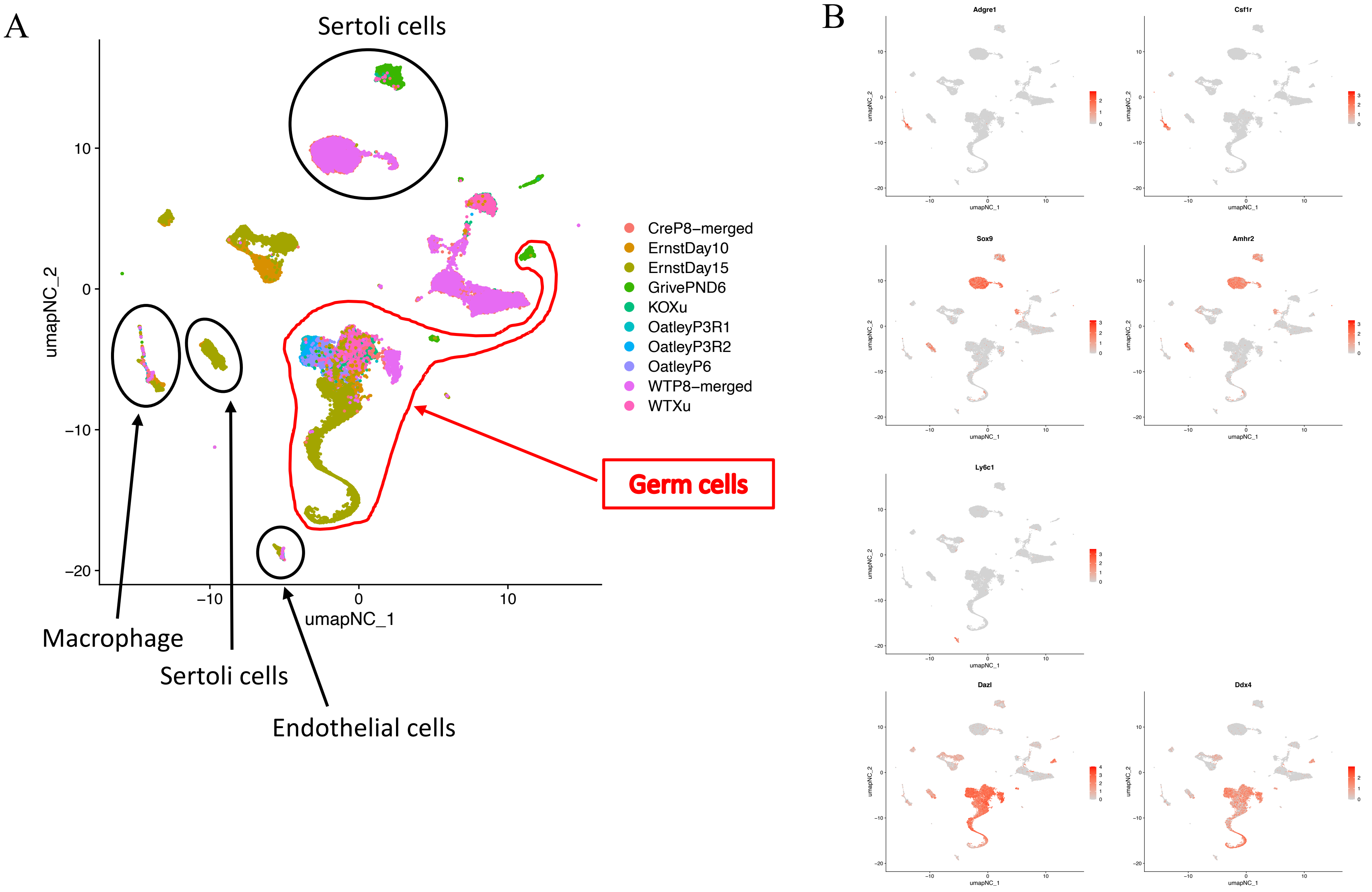

### Figure S4. Identification of the germ cells population based on clustering and marker gene expression

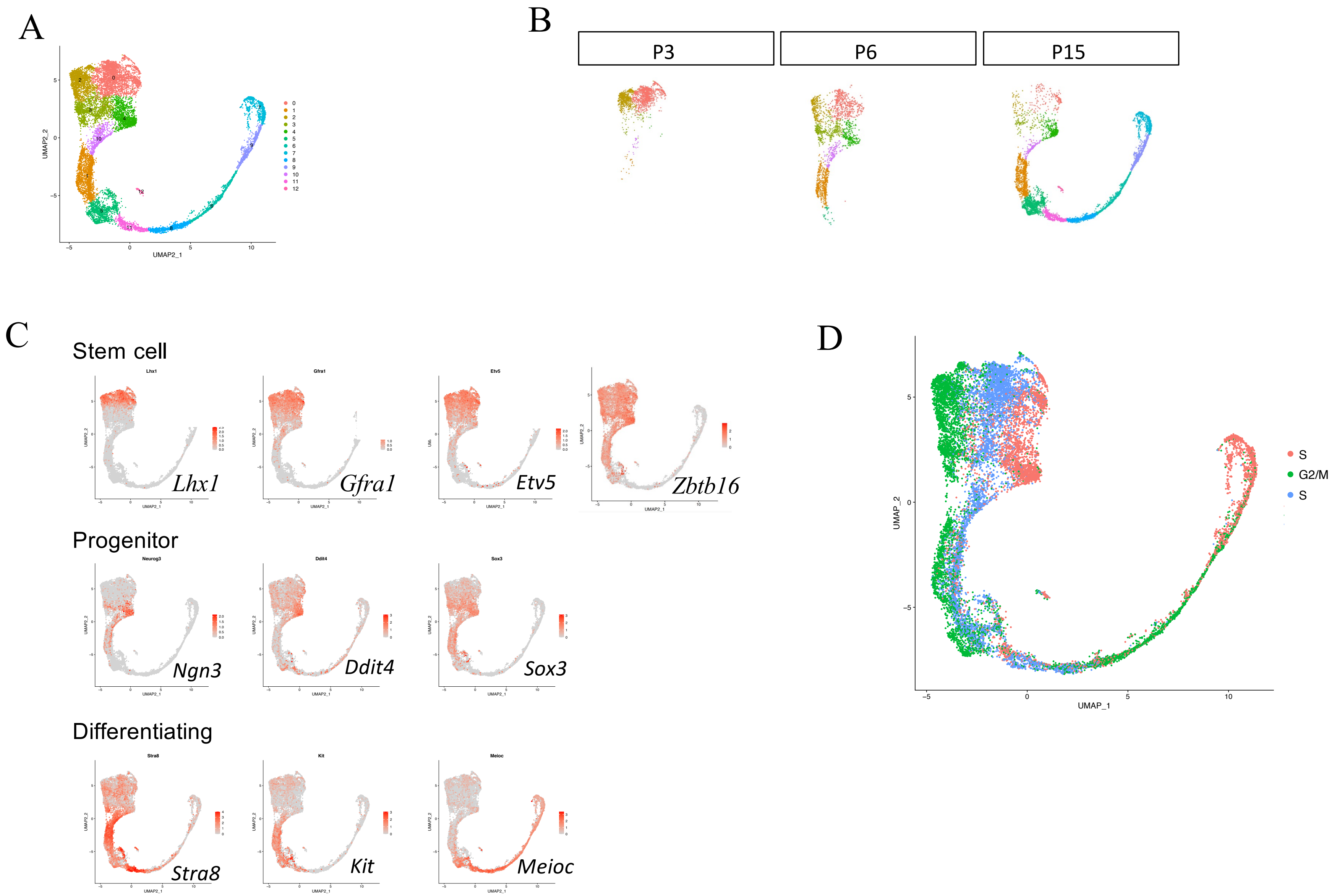

**Figure S5. CNOT3 is required for SSC maintenance and differentiation *in vitro***

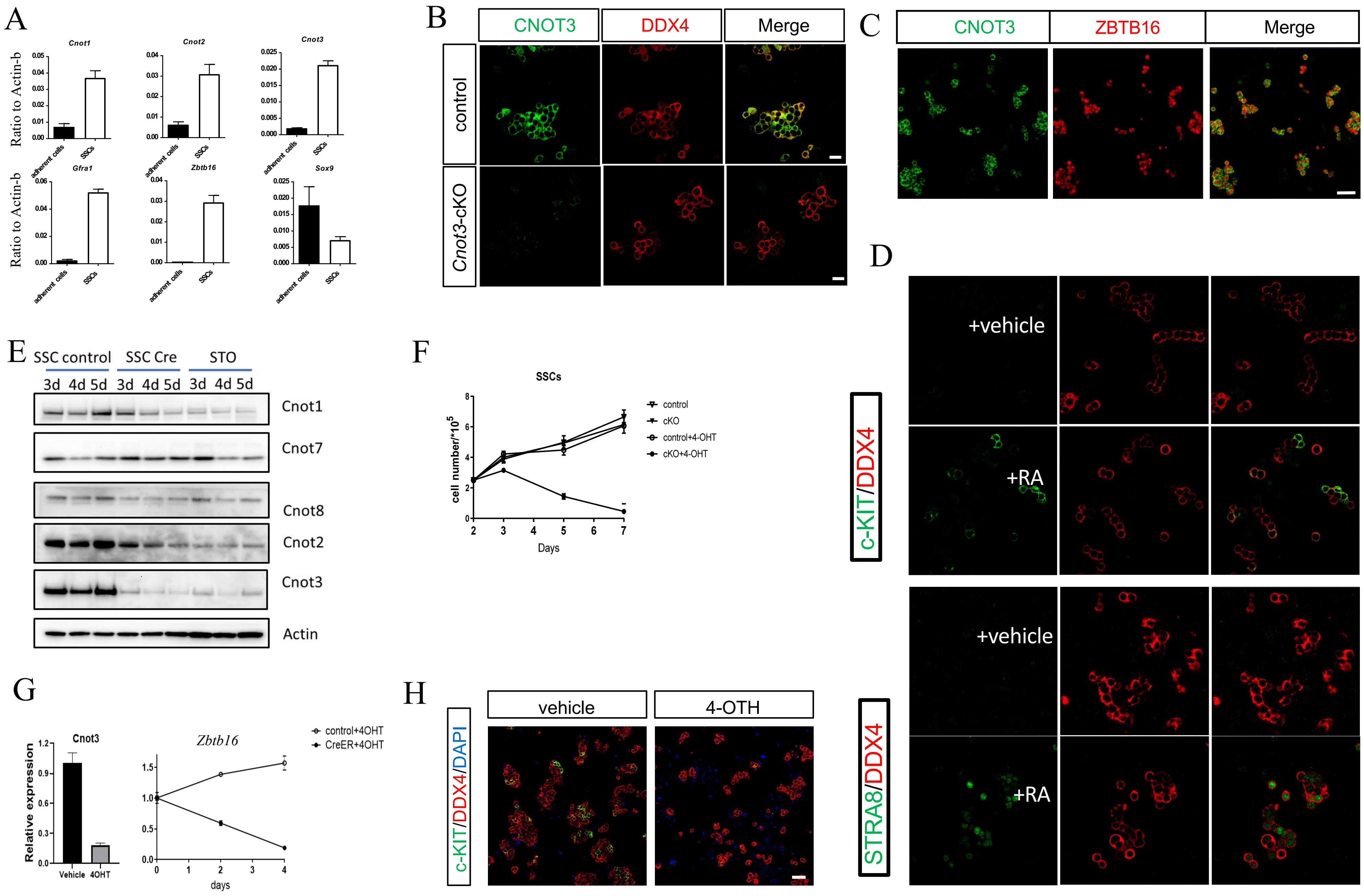

**Figure S6. Effect of *Cnot3* deletion on the glutathione and ROS content in ID4-EGFP+ undifferentiation spermatogonia at P8**

A

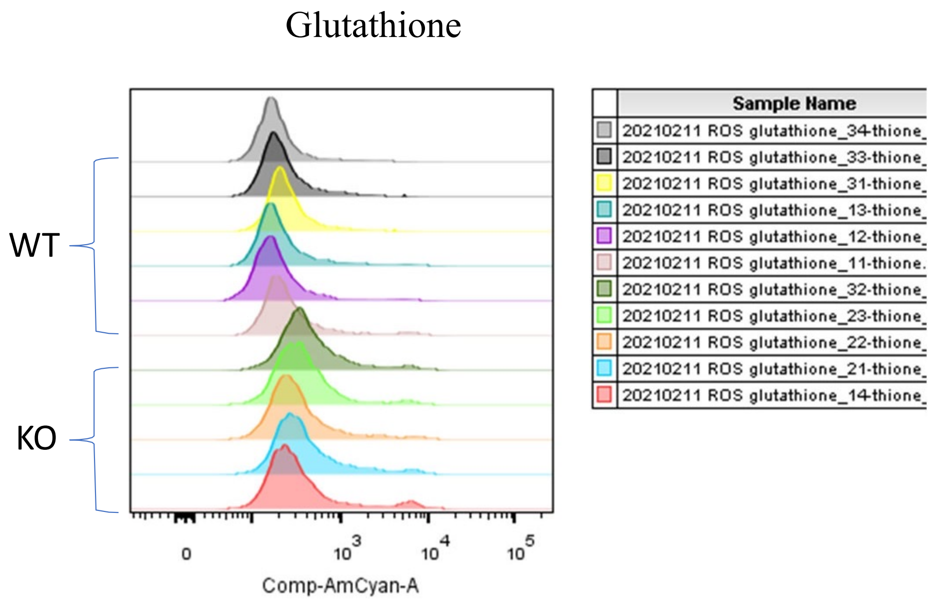


B

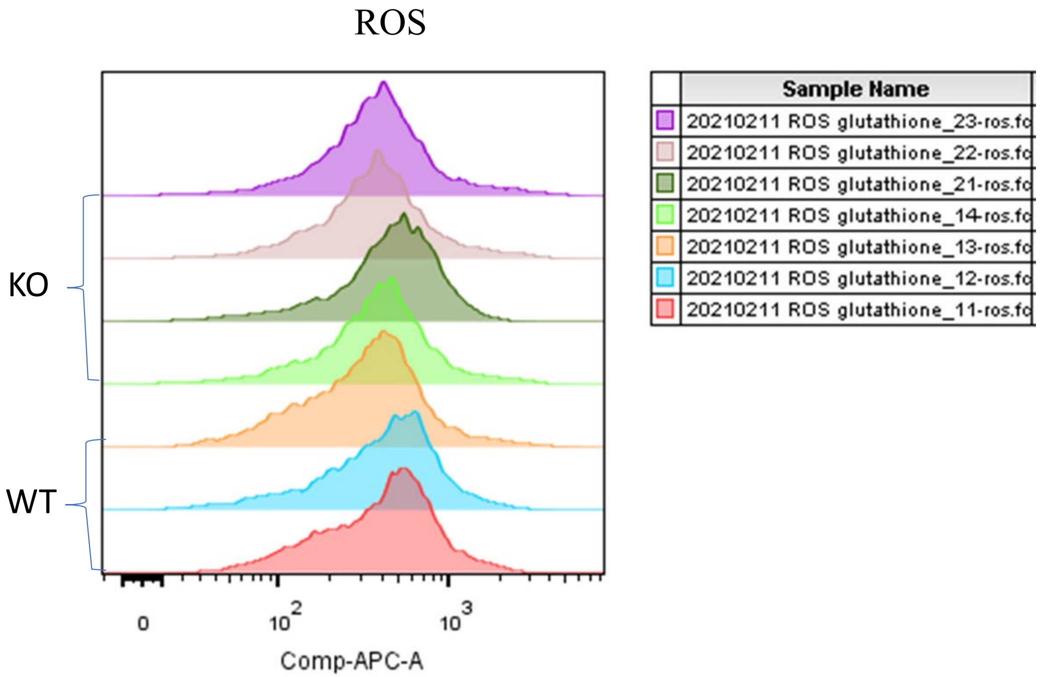
